# Transient Commensal Clonal Interactions Can Drive Tumor Metastasis

**DOI:** 10.1101/2020.01.16.907071

**Authors:** Suha Naffar-Abu Amara, Hendrik J. Kuiken, Laura M. Selfors, Timothy Butler, Marco L. Leung, Cheuk T. Leung, Elaine P. Kuhn, Teodora Kolarova, Carina Hage, Kripa Ganesh, Rosemary Foster, Bo R. Rueda, Athena Aktipis, Paul Spellman, Tan Ince, Nicholas Navin, Gordon B. Mills, Rodrick T. Bronson, Joan S. Brugge

## Abstract

To interrogate functional heterogeneity and crosstalk between tumor cells, we generated clonal populations from a patient-derived ovarian clear cell carcinoma model which forms malignant ascites and solid peritoneal tumors upon intraperitoneal transplantation in mice. The clonal populations were engineered with secreted *Gaussia* luciferase to monitor tumor growth dynamics and tagged with a unique DNA barcode to track their fate in multiclonal mixtures during tumor progression. Only one clone, CL31, grew robustly, generating exclusively malignant ascites. However, multiclonal mixtures formed large solid peritoneal metastases, populated almost entirely by CL31, suggesting that transient cooperative interclonal interactions were sufficient to promote metastasis of CL31. CL31 uniquely harbored *ERBB2* amplification, and its acquired metastatic trait was dependent on transient exposure to amphiregulin, which was exclusively secreted by non-tumorigenic clones. Amphiregulin enhanced CL31 mesothelial clearance, a prerequisite for metastasis. These findings demonstrate that transient, ostensibly innocuous tumor subpopulations can promote metastases via “hit-and-run” commensal interactions.

## Introduction

Tumors consist of diverse subpopulations of neoplastic cells which contribute to intratumoral heterogeneity ^1^. Studies have suggested that genetically and epigenetically heterogeneous subpopulations possess diverse functional abilities ^2, 3^, however, the consequences of this intratumoral functional diversity on behavior of the tumor as a whole is poorly understood. The classical Darwinian model of tumor evolution posits that genetic variants generated during tumor progression compete and, in time, progressively displace one another ^4–6^. However, the long-term maintenance of co-existing genetically distinct subpopulations within the same tumor suggests that cooperation between distinct subpopulations likely influences the tumor phenotype – that is, where one subpopulation provides an essential biological function that is required by others within the tumor, and thus influences the overall composition and behavior of the tumor.

Several studies in mouse models and in *Drosophila* have demonstrated that subpopulations of cells can cooperate to induce tumor growth ^7–9^ and metastasis ^10–15^. In diffuse intrinsic pontine glioma, Vinci and colleagues ^16^ identified a cooperative mechanism between H4K20 methyltransferase-wild-type and -mutant subpopulations that promotes invasion. In all of the aforementioned studies, either specific tumor subpopulations with pre-defined markers or genetically-engineered subclonal populations were examined. Functional studies of intratumoral cooperation during tumor progression using a collection of patient-derived clonal populations without bias toward a specific marker has not been reported.

Multiple studies have tracked clonal populations in the context of tumor progression. Kerso and colleagues ^17, 18^ examined the fate of lentiviral-tagged populations of colon tumor cells during tumorigenesis and demonstrated that the representation of clonal populations changes over time. Using genetic lineage tracing, Driessens et al. ^19^ identified two distinct groups of clones with different proliferation and renewal potential, providing experimental evidence for the existence of cancer stem cells in unperturbed solid tumor growth. However, it was not feasible to address the mechanisms underlying the observed clonal dynamics described in these reports since the clones could not be isolated for mechanistic studies.

Research suggests that cooperative interactions among tumor cells may have important implications for metastasis. For example, Aceto and colleagues discovered that circulating clusters of multiclonal tumor cells were more effective at metastasizing than single circulating tumor cells in a mouse model, and that these clusters were more resistant to apoptosis than single cells ^20^. They also demonstrated that, in patients, higher levels of cell adhesion molecules (plakoglobins) were associated with poorer outcomes. Chapman and colleagues similarly discovered that multiclonal tumor cell groups produce extracellular matrix components and proteases that are associated with greater invasiveness ^21^. These results suggest that cooperation among cancer cells is likely important during invasion and metastasis, but leaves many open questions about the potential mechanisms of molecular crosstalk that underlie this cooperation and how they change over time.

Here, we describe the generation of a collection of single-cell clonal populations from a patient-derived clear cell carcinoma cell line, OCI-C5x ^22^. We then selected a panel of 11 clones based on morphological heterogeneity in culture, and tracked their growth dynamics *in vivo* by assessing Gaussia luciferase activity in blood and characterized the tumorigenicity of each individual clonal population alone or in multiclonal mixtures. By ‘tagging’ each clonal population with a unique DNA barcode, we monitored the clonal dynamics within tumors derived from multiclonal mixtures ^18^. Our findings identify a commensal mechanism of clonal cooperation involving a transient interclonal interaction that promotes metastasis of one clonal population without benefiting the other.

## Results

### Isolation of single-cell clonal populations from a patient-derived clear cell carcinoma model

To investigate whether subpopulations within an individual tumor cooperate to affect tumor behavior as a whole we sought to systematically characterize the tumorigenicity and clonal growth dynamics of individual as well as mixtures of tumor subpopulations *in vivo*. To that end, we isolated single-cell clonal populations from OCI-C5x, an ovarian clear cell carcinoma (CCC) cell line generated from a patient-derived xenograft (PDX) of a treatment-naïve patient primary tumor. OCI-C5x line maintains the spectrum of genetic alterations of the original patient tumor and, when injected into the peritoneum of immunocompromised mice, generates tumors that recapitulate the histopathological features and molecular markers of CCC and of the original tumor ^22^.

To facilitate functional studies and monitoring of tumor burden in mice over time, OCI-C5x cells were engineered to co-express the fluorescent marker tdTomato and Gaussia luciferase (Gluc), a secreted form of luciferase that makes it feasible to track tumor burden by assaying whole blood luciferase activity ^23^. Single OCI-C5x cells were sorted into wells of a 384-well plate and cultured with parental OCI-C5x-condition media to support their initial expansion (Figure S1A). The plates were inspected under a fluorescence microscope four hours after sorting and wells containing more than one cell were flagged and excluded from further analysis. 43 of 90 single cells expanded to generate clonal populations. From these, we chose 11 clones (CL09, CL11, CL12, CL16, CL17, CL28, CL31, CL41, CL44, CL46, and CL49) based on varied morphology in culture for further characterization (Figure S1B). All clonal populations were biobanked as low passage stocks for all subsequent studies.

### Clonal populations exhibit variable growth dynamics *in vivo*

To assess the ability of each individual clonal population to grow after intraperitoneal implantation into immunocompromised mice (referred to as tumorigenicity in this system) and compare it to that of the OCI-C5x parental line or of a heterogeneous mixture of the clonal populations, we injected equal cell numbers (3×10^6^) of each individual clonal population, the OCI-C5x line, or a defined heterogeneous mixture consisting of all 11 clones in equal proportion (referred to as multiclonal mixture) into the peritoneum of immunocompromised female mice (NOD *scid* gamma-NSG) (Figure 1A). Tumor burden was quantified over time via assessment of Gluc activity in the blood ^23^ (Figure 1B). We confirmed that the Gluc levels correlate with cell number *in vivo* in this model (R^2^=0.9949, *P*<0.05) (Figure S2). We found that although all the populations exhibited a dramatic cell loss in the first week following injection, their recovery and growth rate varied (Figures 1B and 1C). Both CL31 and the multiclonal mixture displayed recovery and growth rates that were comparable to the OCI-C5x parental line. CL49 persisted with a lower tumor burden than the parental line and only moderately grew during the duration of the experiment. CL09, CL11, CL12, CL16, CL28, CL41 and CL44 persisted with minimal viability following the initial period of cell loss, but failed to increase in cell number within the experimental time frame. The two remaining clonal populations (CL17 and CL46) exhibited continuous cell loss over time and the Gluc levels in blood were at background levels around the five-week time point on average (Figure 1B). These results indicate that, with the exception of CL31, and to a lesser extent CL49, the majority of the clonal populations lacked tumor forming activity. These results were reproducible across multiple independent experiments performed in a span of over two years, suggesting that this phenotypic diversity is driven by stable cell autonomous properties.

**Figure 1.**
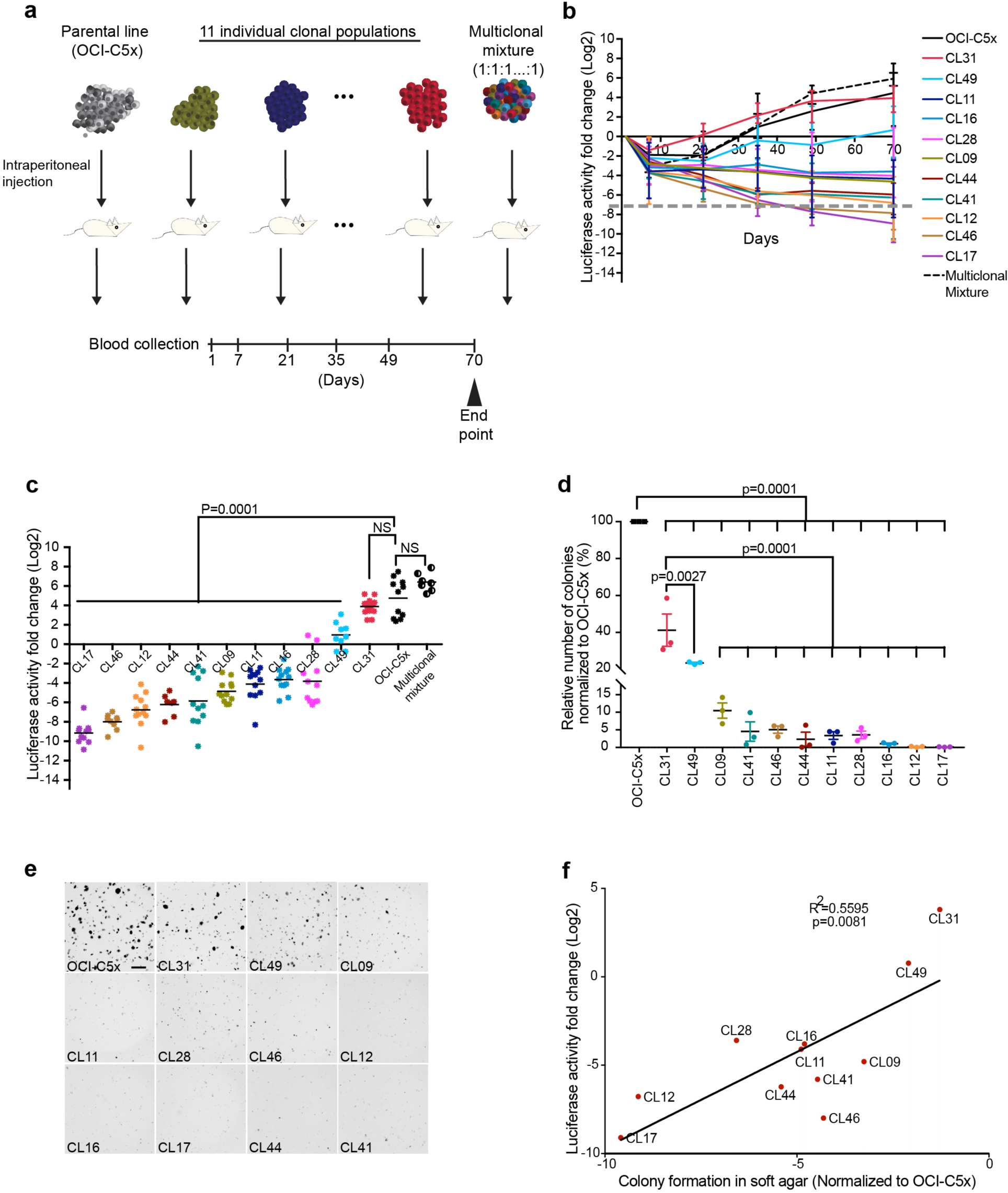
Clonal populations display variable growth dynamics *in vivo*. (A) Schematic depiction of tumor cell populations inoculated into immunocompromised (NSG) female mice. (B) Tumor growth dynamics *in vivo* assessed by measurement of luciferase activity in whole blood samples collected at the indicated time points. Dashed line represents Gluc values at background levels. Data presented as fold change in luciferase activity compared to 24 hours post-injection. (C) Fold change in luciferase activity per mouse determined as in (B) at the 10 week end point. *P* values were computed using the one-way ANOVA test followed by Dunnett’s multiple comparison test, comparing parental line to each of the clones and the multiclonal mixture. NS: not significant. (D) Colony formation in soft agar measured after 21 days and normalized to the parental line. Data represent mean ± SEM from three independent experiments, biological replicates were performed on cells from that had been passaged anywhere between eight-23 times. *P* values were computed using the one-way ANOVA test followed by Dunnett’s multiple comparison test, comparing either parental or CL31 against each of the clones. (E) Images of a representative soft agar colony formation assay. Scale bar represents 1mm. (F) Linear regression analysis of colony formation in soft agar and tumor burden (measured by luciferase activity in blood samples collected at 10 week end point) for all 11 clonal populations. See also Figures S1, S2 and S3.

To investigate whether the tumorigenicity of the clonal populations could be attributed to enhanced proliferative capacity, we assessed the correlation between *in vitro* doubling time in monolayer culture and tumor growth (Gluc levels) (Figure S3A-S3B). We found that tumorigenicity did not correlate with *in vitro* proliferation rates (R^2^=0.1248, *P*=0.3763). Next, we examined the ability of the clonal populations to form colonies in soft agar, a commonly used surrogate for tumor formation. Among the clonal populations, CL31 formed the most colonies followed by CL49, and the remaining clones formed few, if any, colonies. This *in vitro* phenotypic diversity was stable across three independent experiments, performed with cells passaged between eight to 23 times. Tumorigenicity correlated with colony formation in soft agar (R^2^=0.5595, *P*=0.0081) (Figure 1D-1F), suggesting that anchorage independence reflects tumorigenicity in our model system.

### Individual clonal populations fail to form solid peritoneal metastases

Ovarian carcinoma metastatic dissemination is distinct from that of other solid cancers in that it is primarily non-hematogenous and limited to the peritoneal cavity, where tumor cells, once exfoliated from the primary site, are spread throughout the abdominal cavity and grow in the peritoneal fluid as small clusters (malignant ascites) and form solid peritoneal masses on the mesothelial surfaces, with clear preferences for the omentum and the diaphragm ^24^. The pattern of growth of the OCI-C5x cell line in immunocompromised mice recapitulates these disease features, consistently generating the two forms of tumor cell growth: malignant ascites, with 400-1000μl of cell pellet volume (Figure 2A), as well as solid peritoneal metastases on the mesentery, ovary, and large tumors on the diaphragm that often extend to the liver (Figure 2B).

**Figure 2.**
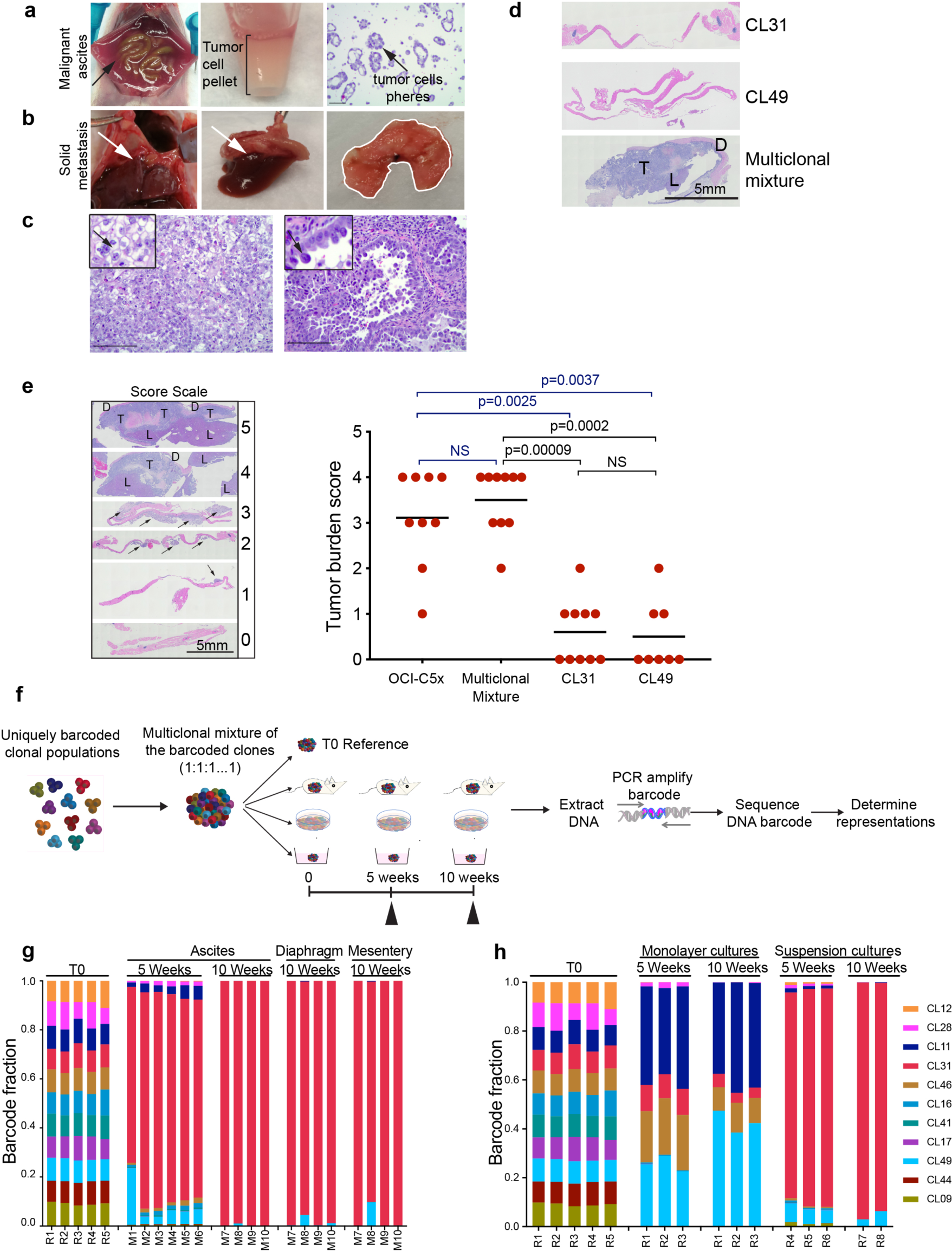
Individual clonal populations fail to form solid peritoneal metastases. (A) Representative images of OCI-C5x generated malignant ascites, and (B) solid tumors on the diaphragm, which connect to the liver (white arrow) (C) H&E-stained section from the solid tumor masses. Note that the tumor cells maintain histopathological characteristics of CCC in patients with “clear” cells (C-left image, arrow) and hobnail features (C-right image, arrow). Scale bar, 100μm. (D) Representative H&E images of tumor sections of solid metastases on diaphragm generated by CL31, CL49 and the multiclonal mixture. T: tumor cells. D: diaphragm. L: liver. Scale bar, 5mm. (E) Tumor burden score of solid metastases on the diaphragm (Data pooled from two independent experiments, n=8-10 mice per group). Scored blindly. *P* values were computed using the Chi-square test with Monte Carlo simulation. Schematic depiction of experimental design and workflow for barcode experiments. Barcode representation in the indicated samples collected from mice injected with the multiclonal mixture composed of barcoded clones, and (H) in samples collected from monolayer and suspension cultures *in vitro*. A representative of two independent experiments.

Moreover, OCI-C5x tumors exhibit “clear” and “hobnail” cells, typical CCC histopathological features (Figure 2C). Consistent with the blood Gluc assay results, only CL31, CL49, and the multiclonal mixture produced any detectable tumor cell growth in immunocompromised mice. CL49 displayed weak *in vivo* cell growth, generating only 10-50μl cell pellet volume of malignant ascites, whereas CL31, like the OCI-C5x parental line, consistently generated more than 400μl (ranges between 400-1000μl) cell pellet volume of malignant ascites. However, unlike OCI-C5x, neither CL31 nor any of the other clones generated macroscopically detectable solid peritoneal metastases. In contrast, the multiclonal mixture fully recapitulated the phenotype of the parental line, generating robust malignant ascites (>400μl cell pellet volume) as well as solid peritoneal metastases that were most prominent on the diaphragm and often extended to the liver (Figure 2D). Blind scoring of tumor burden in the diaphragm histology sections from mice injected with CL31, CL49, OCI-C5x, or the multiclonal mixture confirmed that CL31 and CL49 failed to recapitulate the solid peritoneal metastasis phenotype of the OCI-C5x parental line or the multiclonal mixture (Figure 2E). These data strongly suggest that the OCI-C5x-derived clonal populations cooperate to generate solid peritoneal metastases, a phenotype that the individual populations are incapable of producing on their own.

### Malignant ascites and solid peritoneal metastases derived from the multiclonal mixture are dominated by CL31

To identify the clonal representation within the malignant ascites and solid peritoneal metastasis generated by the multiclonal mixture, we tagged each clonal population with a unique semi-random 30 base-pair DNA barcode with balanced GC content (50%) to ensure uniform PCR-amplification efficiency across the barcodes ^18^. To spatially and temporally detect the representation of each clonal population during tumor progression *in vivo*, malignant ascites were collected at five and ten weeks after IP injection of the multiclonal mixture and solid peritoneal metastasis at 10 weeks. In parallel, we included two *in vitro* arms: (1) standard monolayer culture of the multiclonal mixture passaged every four days, and (2) suspension culture of the multiclonal mixture in which media was refreshed every four days. The genomic DNA of all tumor samples and cell cultures collected at five and 10 weeks was extracted, and the barcode sequences were PCR-amplified and subjected to Next Generation Sequencing (NGS). The barcode representations were compared to that of the initial (T0) multiclonal mixture (Figure 2F). The barcode representation analyses indicated that, at the five-week time point, the malignant ascites consisted of multiple clonal populations, with a clear dominance of CL31 (82.1% on average); other clones (predominantly CL09, CL11, CL12, C28 and CL46) were present at lower frequencies (<30% combined). At the 10-week time point, the tumor samples consisted almost entirely of CL31 (>97% at all sites) (Figure 2G) and the relative abundance of the other clones dropped below 0.0001% at all sites; this represented the “noise threshold cutoff” values of the barcode reads, with exception of CL49 (0.25% in ascites, 1.25% diaphragm, and 2.25% mesentery on average). The presence of the minor clones at the five-week time point was consistent with the tumor forming ability of these specific clones as they maintained minimal viability when injected individually (Figure 1B). Moreover, the dominant representation of CL31 in the malignant ascites at five and ten weeks was in agreement with this clone’s individual, robust tumor forming capacity (Figure 1B). However, the dominance of CL31 in the solid peritoneal tumors on the mesentery and diaphragm was unexpected since no solid peritoneal metastases were detectable when CL31 alone was injected IP (Figure 2E). This dominance of CL31 was not recapitulated in monolayer cultures where CL31 was outcompeted at early time points by CL11, CL28, CL46 and CL49 (Figure 2H), suggesting that the dominance of CL31 *in vivo* is not due to superior proliferative fitness in the clonal mixture. In contrast to the monolayer culture clonal representation, the pattern of clonal dynamics observed under suspension conditions *in vitro* largely recapitulated that observed *in vivo*, with CL31 outcompeting the rest of the clones as early as the five-week time point. These results are in line with the anchorage independence of this clone and are consistent with anchorage independent survival being associated with the competence to grow *in vivo* in ascites fluid. These results, together with the finding that CL31 is limited to generation of malignant ascites and is unable to produce solid peritoneal metastases on its own, provide strong evidence that interclonal cooperation is required to promote the acquisition of solid peritoneal metastasis activity by CL31.

### Genetic analyses identify differential *ERBB2* amplification in CL31

To investigate the molecular mechanism(s) underlying this clonal cooperation, we performed multiple genomic analyses. Copy number aberration (CNA) profiling of the OCI-C5x parental line revealed that OCI-C5x is mainly diploid with no dominant regions of chromosomal deletions or amplification detected in bulk analysis (Figure 3A). However, the genetic heterogeneity of this cell line became apparent when the clonal subpopulations were analyzed. For example, while most of the clones are diploid, CL11, CL44, and CL46 harbor whole chromosome loss (e.g. chr 3, 4, 6, 10, 11, 13 and 16) or gains (7, 12 and X), a phenomenon commonly observed in CCC ^25, 26^. Notably, CNA analysis also revealed a focal amplification of *ERBB2* exclusively in CL31 (the amplified region of chromosome 17 which contains six copies of *ERBB2* is marked with a star) (Figure 3A). Fluorescence in situ hybridization (FISH) confirmed the *ERBB2* amplification in CL31. The majority of the cells (99%) had an *ERBB2*:chromosome 17 (centromere CEP17) ratio of two or more with an overall average ratio of 2.54 (Figure 3B).

**Figure 3.**
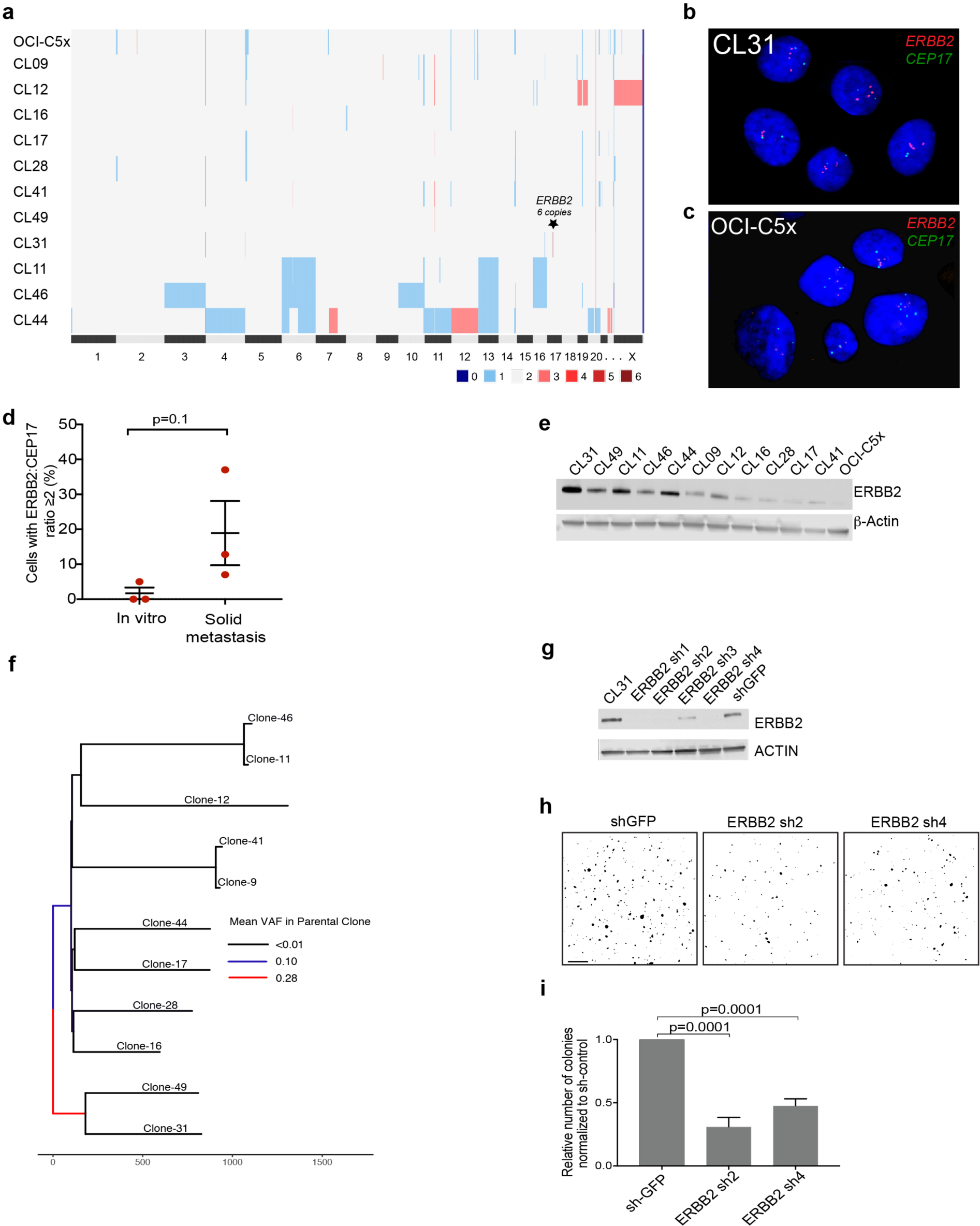
Molecular analyses of clonal populations reveal CL31-specific genetic alterations. (A) Heatmap representation of copy number alterations in the parental OCI-C5x cell line and clonal populations. (B-C) Representative images of FISH staining for *ERBB2* (red) and CEP17 (green) of (B) CL31 and (C) parental OCI-C5x cells *in vitro*. (D) Quantification of cells with *ERBB2:CEP17* copy number ratio equal or greater than two (FISH data) in the OCI-C5x cell line *in vitro* from experiment in (B) and in solid metastases derived from the OCI-C5x cell line. Data shown as mean ± SEM from three independent experiments, *P* value was computed using the Mann-Whitney test. (E) Representative Western blot of ERBB2 protein in the OCI-C5x cell line and each of the clonal populations. β-actin was used as a loading control. (F) Maximum parsimony tree generated from all filter-passing substitutions detected by whole-exome sequencing of all single-cell derived clonal populations, and an autologous DNA sample from the patient’s blood. Branches colored by mean VAF of mutations in parental clone. (G) Western blot showing *ERBB2* knockdown in CL31 using four distinct shRNAs. (H) Representative images and (I) quantification of colony formation in soft agar by CL31 expressing a control shRNA (GFP) or one of two distinct *ERBB2* shRNAs (Sh2 and Sh4). Scale bar, 500μm. The number of colonies were normalized to that of shGFP control. Three independent experiments were summarized by mean ± SEM. *P* values were computed using the one-way ANOVA test and corrected for multiple comparisons using Dunnett’s method. See also Figures S4 and S5.

To test whether *ERBB2* amplification pre-existed in the original patient sample from which OCI-C5x was derived, we performed FISH analysis on histology section of the original primary tumor (ovary primary site). The sample was scored twice by two independent technicians at the Cytogenomics Core Laboratory (Brigham and Women’s Hospital, Boston, MA) for *ERBB2*:CEP17 ratio in ten different regions identified visually by a pathologist as tumor cell-enriched areas and confirmed by staining with S100A1, a marker that differentially identifies ovarian carcinoma cells from normal ovarian tissue ^27^ (Figure S4A). In three regions (regions #3, #4 and #6), more than 20% (26%, 23.3% and 25% respectively) of cells displayed *ERBB2*:CEP17 signal ratio equal or greater than two (cells with only 1 CEP17 signal were excluded), with an average ratio of *ERBB2*:CEP17 signal of 2.96, 2.42 and 2.7 respectively (Figures S4B-S4C and S4E). Immunohistochemistry confirmed that a subset of tumor cells within region #3 express elevated levels of ERBB2 protein (Figure S4D). These results demonstrate that *ERBB2* amplification was present in the original patient tumor and was not acquired during derivation of the OCI-5Cx cell line.

We also examined the abundance of *ERBB2* amplified cells in the OCI-C5x parental line by FISH analysis. In one out of three experiments, 5% of 100 scored cells displayed an *ERBB2*:CEP17 signal ratio equal to two; in two additional experiments in which 100 cells were scored in each, none displayed an *ERBB2*:CEP17 signal equal or greater than two. These results indicate that cells with low level amplification of *ERBB2* are rare in the parental cell line. Conversely, solid peritoneal metastases generated by the parental line contained a higher percentage of cells with an *ERBB2*:CEP17 ratio equal or greater than two (12%, 37% and 7% in three independent xenografts), with average *ERBB2*:CEP17 ratio of 2.52±0.15 (Figure 3D). While not statistically significant (p=0.1), this shows a trend towards enrichment of cells with extra copies of *ERBB2* in the metastatic tumors, suggesting selection *in vivo*. Moreover, analysis of ERBB2 protein levels by Western blotting confirmed that, among all the clones and the OCI-C5x parental line, CL31 has the highest levels of this protein while the OCI-C5x parental line has the lowest levels (Figure 3E). Taken together, these analyses revealed alterations that would have otherwise been masked in bulk analysis of the OCI-C5x population and indicate that the *ERBB2* amplification is present at a low frequency in the original tumor and the OCI-C5x parental line derived from it *in vitro*.

We also performed whole exome sequencing of all clones, the OCI-C5x parental line, and an autologous DNA sample from the patient’s blood, which revealed an extremely high mutation burden (106 mutations per megabase) that is likely driven by mutations in the mismatch DNA repair gene *MSH2* and the DNA damage response gene *ATM*, shared across all samples. This is also supported by the finding that over half of mutations can be attributed to mutational signatures 6 and 20 which have been previously associated with mismatch repair gene defects ^28^ (Figure S5A-S5B). In addition, we found no significant variation in the proportion of signatures present in any of the clones. Maximum parsimony-based phylogenetic analysis of whole-exome mutations indicated that CL31 and CL49, the only tumor forming clones, were located on one branch of the maximum parsimony tree, suggesting that they are more closely related genetically to each other than to other clones (Figure 3F). Indeed, although all of the clones are related, CL31 and CL49 are quite distant from the other clones on the parsimony tree. We also analyzed single nucleotide variation (SNV) allele frequencies of mutations belonging to each branch of the tree in the OCI-C5x parental population.

This revealed that the mutations shared by CL31 and CL49 are at a higher abundance in the parent than any other clones (Wilcoxon, p<0.0001). This finding suggests that a large proportion of the OCI-C5x parental line is comprised of a common ancestor of these two clones. Moreover, these data imply that *ERBB2* amplification, which was present only in CL31, may be a later event acquired in a small subset of the original tumor population.

### Elevated ERBB2 levels enhance anchorage independent growth of CL31

The *ERBB2* amplification in CL31 was of particular interest because this clone is the only robust tumor-forming clone. Moreover, *ERBB2* amplification and overexpression are more common in CCC than in other ovarian cancer subtypes ^29–31^. To examine whether the elevated ERBB2 levels in CL31 contribute to its robust phenotype, we evaluated the effects of shRNA-mediated *ERBB2* downregulation on soft agar colony formation, which correlated with tumorigenic ability *in vivo* in our model (Figure 1F). Two distinct hairpins that induced dramatic reduction in ERBB2 protein levels impaired CL31 colony formation by approximately 50% compared to the sh-GFP control (Figures 3G-3I). Together, these results suggest that elevated ERBB2 protein levels enhance anchorage independent growth of CL31 and, since anchorage independence correlated with tumor growth ability *in vivo* in our system, suggests that the acquisition of anchorage independence contributes to the tumor-forming activity of CL31.

### Overexpression of amphiregulin induces peritoneal metastasis of CL31

We next investigated whether CL31’s new acquired metastatic phenotype is due to secreted factors that are either lowly-or non-expressed in CL31 and supplied by one or more of the other clones. To identify such factors, we performed RNA-seq analysis and filtered the candidate secreted factors for those whose corresponding receptor(s) were expressed in CL31 (Figure 4A). We identified five candidate factors, three of which were EGFR family ligands [amphiregulin (AREG), Betacellulin (BTC), and Epiregulin (EREG)] that activate ERBB2. AREG, in particular, was an attractive target not only because it was the most statistically significant factor identified by RNA-seq, but also because previous studies have shown that AREG is present at elevated amounts in ascites fluid collected from advanced stage ovarian and lung cancer patients ^32^, its expression has been associated with poor prognosis in several cancer types ^33^, and it has been implicated in invasion of ovarian cancer cells ^34, 35^.

**Figure 4.**
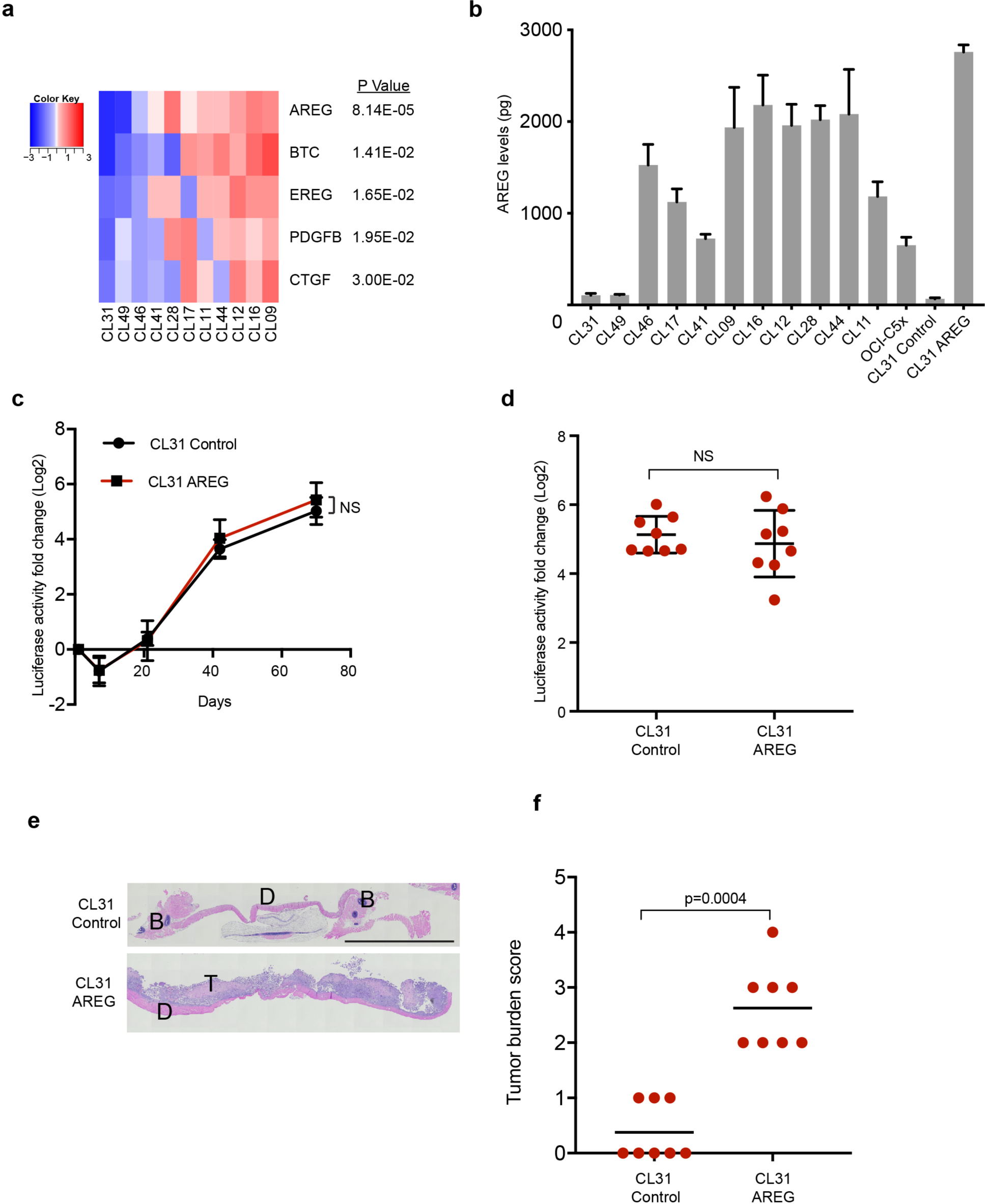
Overexpression of amphiregulin induces peritoneal metastasis of CL31. (A) Heatmap of ligands identified by RNA-sequencing. Growth factors and cytokines that were expressed by one or more of the other clones and poorly expressed in CL31 were identified (FDR-corrected p<0.05, edgeR) and filtered for those whose corresponding receptor(s) were expressed in CL31 (cpm>2). The relative mRNA levels of the identified ligands/growth factors are shown. Data are mean centered Log2. (B) Levels of AREG secreted into the media by the indicated clonal populations over the course of 36 hours, as determined by an ELISA assay. Data represent as mean ± SEM from three independent experiments. (C) Tumor burden *in vivo* assessed by measurement of luciferase activity in whole blood samples collected at the indicated time points. Data presented as fold change in luciferase activity compared to 24 hours post-injection. Representative of two independent experiments. Data summarized by mean ± SD, *p* values were computed using the Student’s t test and FDR corrected. (D) Fold change in luciferase activity in blood samples of individual mice collected at the endpoint (10 weeks) relative to the 24h time point. The data shown as mean ± SD. *p* values were computed using the one-way ANOVA test and Dunnett’s multiple comparison test. (E) Malignant ascites cell pellet volume at the endpoint (10 weeks). The data shown as mean ± SD. *p* values were computed using the Mann-Whitney test. NS: not significant. (F) Representative H&E images of solid peritoneal metastases on the diaphragm of mice inoculated with CL31 expressing AREG or vector control. T: tumor cells. D: diaphragm. Scale bar, 2mm. (G) Tumor burden score of solid metastases on the diaphragms of mice inoculated with CL31 expressing AREG or vector control, determined as in Figure 2E. *p* value was computed using the Chi-square test with Monte Carlo simulation. (D-G) Data pooled from two independent experiments, total n=8 mice. See also Figure S6.

To test whether AREG is sufficient to induce solid metastases formation by CL31, we engineered this clone to exogenously express AREG at a level comparable to that observed in the other clonal populations (Figure 4B and Figure S6A) and examined its aggressiveness and metastatic ability in two independent experiments. AREG overexpression in CL31 did not affect its tumor growth dynamics *in vivo* (based on Gluc activity in blood) (Figure 4C-4D), nor the cells doubling rate *in vitro* (Figure S6B).

However, AREG overexpression in CL31 did induce the formation of solid peritoneal metastases on the diaphragm (Figure 4E and 4F). This result indicates that AREG overexpression is sufficient to induce a phenotypic switch in CL31 from malignant ascites to solid metastatic phenotype.

### AREG acts during an early temporal window to induce solid peritoneal metastasis

The barcode analyses of the tumors generated from the multiclonal mixture as well as the growth dynamics of the individual clones (Figures 1B and 2G) indicate that the representation of AREG-high clones is dramatically lower at three-to five-weeks after implantation *in vivo*, suggesting that if AREG contributes to the solid peritoneal metastasis of CL31 within the multiclonal mixture, it likely acts during an early temporal window to induce the phenotypic switch in CL31 and is later dispensable. To test this hypothesis, we examined whether brief supplementation with recombinant human AREG during the three weeks following injection of CL31 alone could induce the metastatic phenotype. In an attempt to mimic the gradual decrease of AREG provided by the clones *in vivo* (due to continuous cell death), mice were injected intraperitoneally with either recombinant human AREG (5μg/mouse) or vehicle control over the course of the first three weeks as follows: daily administration starting on day 1 during the first week, then every other day in the following week, and twice a week during the third week (Figure 5A). Supplementation with recombinant human AREG significantly increased the formation of solid peritoneal metastases by CL31 (Figure 5B and 5C) compared to vehicle control without significantly affecting tumor cell growth rate (Figure 5D). This data suggests that AREG availability during the initial and brief temporal window is sufficient to induce the phenotypic switch in CL31 to produce solid peritoneal metastasis and that this phenotype is not due to increased tumor growth *in vivo*. This data also supports our hypothesis that the transient presence of AREG-high non-tumorigenic clones may be sufficient to induce this phenotypic switch in CL31 *in vivo*.

**Figure 5.**
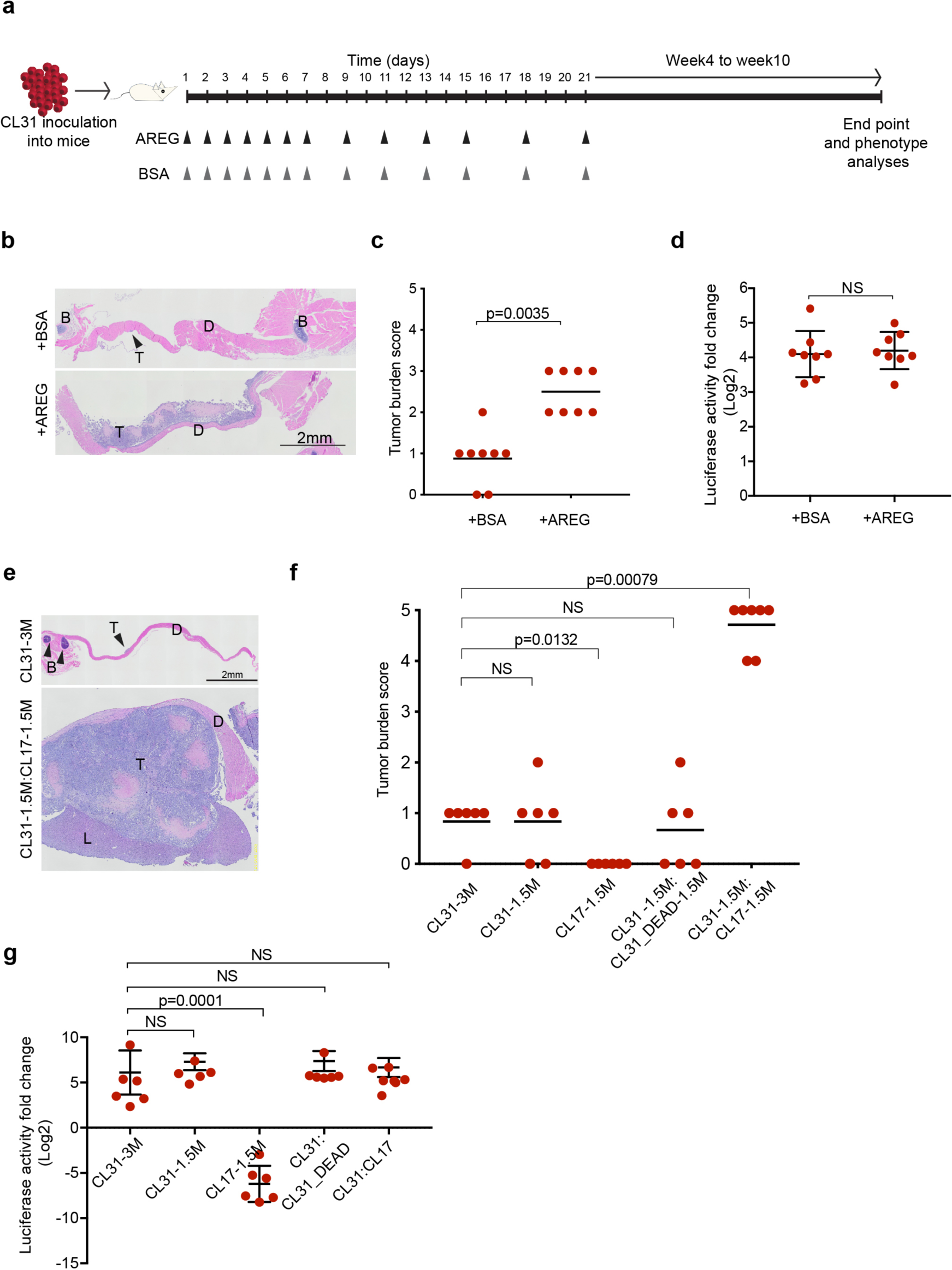
Transient exposure to CL17 or AREG is sufficient to promote peritoneal metastasis of CL31. (A) Schematic depiction of experimental design of short-term supplementation with human recombinant AREG *in vivo.* (B) Representative H&E images of diaphragms collected from mice at the 10-week endpoint supplemented with AREG or BSA vehicle control as depicted in (A). T: tumor cells. D: diaphragm. B: Bone. Scale bar, 2mm. (C) Tumor burden score of solid metastases on the diaphragm generated by CL31 in mice treated with AREG or BSA vehicle control. Data pooled from two independent experiments, total n=8 mice. *p* value was computed using the Chi-square test with Monte Carlo simulation. (D) Fold change in luciferase activity in blood samples collected from individual mice at the endpoint (10 weeks) relative to the 24-hour time point. Data pooled from two independent experiments (total n=8 mice) and shown as mean ± SD. *p* value was computed using the Mann-Whitney test. NS: not significant. (E) Representative H&E images of the diaphragms of mice inoculated with CL31 alone (3×10^6^ cells) or a 1:1 mixture of CL31 and CL17 (3×10^6^ total cells). T: tumor cells. D: diaphragm. L: liver. B: Bone. Scale bar, 2mm. (F) Tumor burden score of solid metastases on the diaphragm generated by the indicated cell inocula. Data pooled from two independent experiments (total n=6-7 mice) and shown as mean ± SD. *p* values were computed using the Chi-square test with Monte Carlo simulation. NS: not significant. (G) Fold change in luciferase activity in blood samples collected from individual mice at the endpoint (10 weeks) relative to the 24-hour time point. Data pooled from two independent experiments (total n=6-7 mice) and shown as mean ± SD. *p* values were computed using the one-way ANOVA test and Dunnett’s multiple comparison test. NS: not significant.

### Transient and non-tumorigenic AREG-high clone promotes CL31 phenotypic switch

To further examine whether the transient presence of a non-tumorigenic AREG-high clone is sufficient to induce the phenotype switch in CL31, we assessed the phenotype of a pairwise combination of CL31 with CL17, an AREG-high non-tumorigenic clone. CL17 was a particularly ideal clone for this experiment since it is the only AREG-high clone that continuously decreases in cell number throughout the experimental time frame and its Gluc levels in blood reached background levels at around 5 weeks (Figure 1B), suggesting that this clone is short-lived. We found that a 1:1 mixture of CL31 and CL17 (3 ×10^6^ total cells), but not CL31 alone (either 1.5×10^6^ or 3×10^6^ cells), generated large solid peritoneal metastases, most prominently on the diaphragm with extension to the liver (Figure 5E). This result was not due to secondary effects induced by rapid death of CL17 (e.g. due to an inflammatory response), as a 1:1 mixture of live and dead CL31 cells (generated by repeated freeze-thaw) failed to recapitulate the metastatic phenotype of the pairwise CL31:CL17 mixture (Figure 5F).

Notably, the overall tumor burden (based on blood Gluc levels) of mice injected with CL31:CL17 was comparable to that of CL31 alone (Figure 5G), indicating that the metastatic phenotype is not due to increased tumor cell growth. These results suggest that a transiently existing non-tumorigenic clone is sufficient to confer solid metastasis formation ability on CL31.

Next, we addressed whether the ability of CL17 to induce the phenotypic switch in CL31 is dependent on AREG. Following intraperitoneal injection of the CL31:CL17 mixture, mice were treated twice a week with either an AREG blocking antibody or vehicle control, and tumor cell growth was measured over time by quantifying blood Gluc levels. We found that treatment with AREG antibody reduced the overall tumor burden (based on blood Gluc levels, p=0.012) (Figure 6A and 6B); however, there was no significant reduction in the cell pellet volume of malignant ascites (Figure 6C), suggesting that AREG antibody did not reduce the growth of CL31 tumor cells in the ascites. In contrast, the burden of solid metastases on the diaphragm was significantly lower in the AREG antibody-treated group compared to vehicle control (Figure 6D-6F), suggesting that AREG blocking antibody specifically interferes with the formation of solid peritoneal metastases.

**Figure 6.**
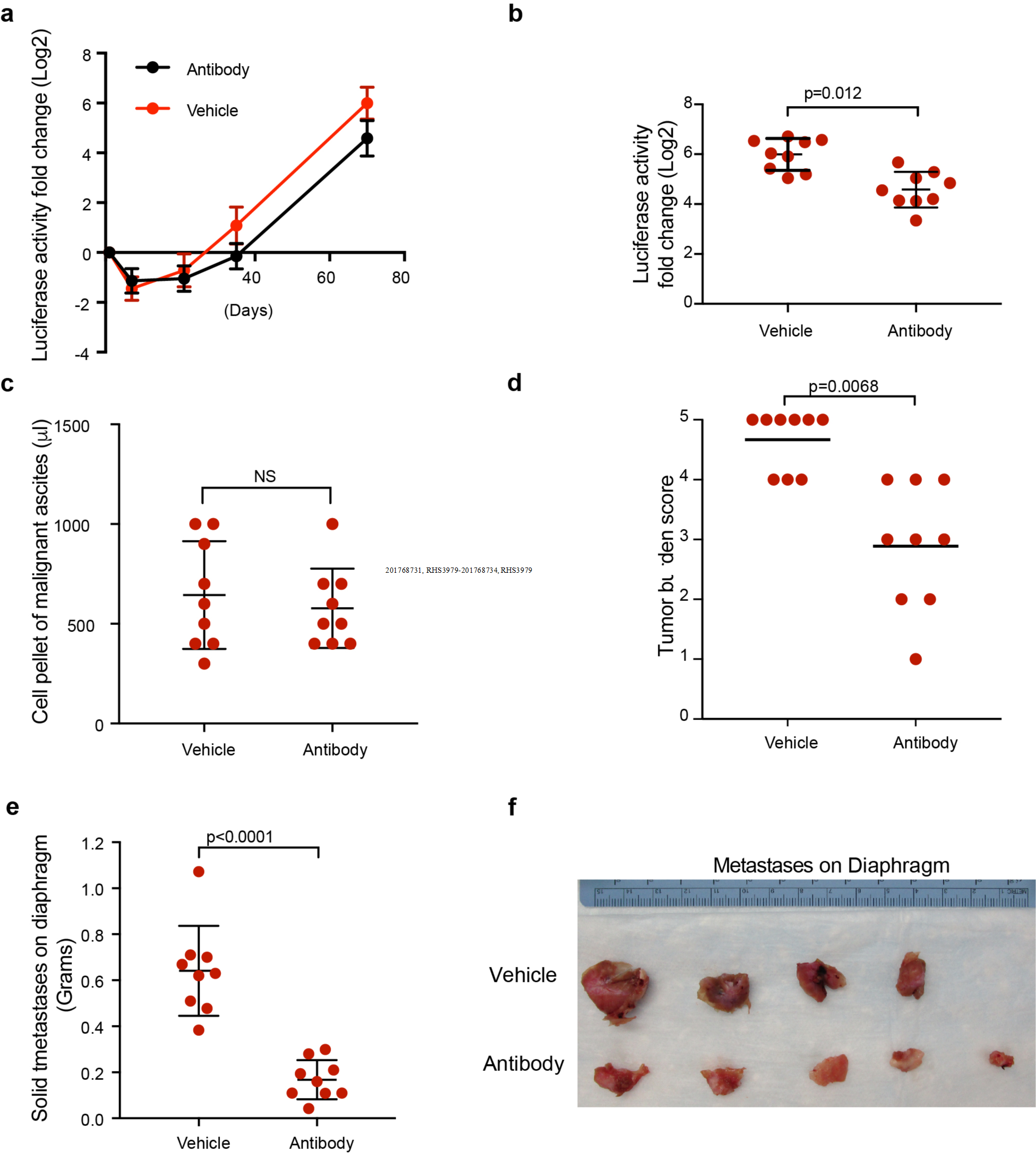
AREG blocking antibody reduces solid peritoneal metastasis. A total of 3×10^6^ cells of an equal mixture of CL31 and CL17 were injected I.P. Mice were treated with 200μg monoclonal AREG blocking antibody immediately following injection and twice a week thereafter. Control group was treated with PBS vehicle control. (A) Tumor growth dynamics *in vivo* assessed by measurement of luciferase activity in whole blood samples collected at the indicated time points. Data presented as fold change in luciferase activity compared to 24 hours post-injection. (B) Fold change in luciferase activity per mouse determined as in (A) at the 10 week endpoint. (C) Malignant ascites cell pellet volume at the 10-week endpoint. (D) Tumor burden score of solid metastases on the diaphragm at 10-week endpoint generated by the mixture of CL31 and CL17 in mice treated with AREG blocking antibody or vehicle control. (E) Weight of solid tumors on the diaphragms at 10-week endpoint of same tumors in (D). Data pooled from two independent experiments, nine mice in total for each group, and shown as mean ± SD. *P* values from Mann-Whitney test. NS: not significant. (F) Individual diaphragms from mice in one of the two experiments at the 10-week endpoint.

To test the abundance of CL31 and CL17 in the resultant tumors of CL31:CL17 mixture, barcode representation analysis was performed on malignant ascites and solid mases on the diaphragm at the 10-week time point. The results showed that CL17 barcode reads were below background levels (defined as the barcode count for a CL11, which was not present in the mixture, was used as a reference to set the noise threshold of the barcode counts) suggesting that CL17 does not co-exist with CL31 in the solid peritoneal metastases (Figure S7).

Together, the results from the above experiments involving multiple approaches to evaluate the role of non-tumorigenic clones and AREG in regulating CL31 metastasis provide strong evidence that transient and non-tumorigenic clones cooperate with CL31 to affect tumor aggressiveness and that the interclonal cooperation is, at least in part, dependent on AREG.

### AREG enhances mesothelial clearance

Dissemination of ovarian carcinoma cells to distant sites involves attachment to and clearance of the superficial layer of the mesothelium that encloses the organs in the peritoneal cavity ^36–38^. To investigate whether the induction of CL31 metastatic ability by AREG is due to enhanced mesothelial clearance, we utilized a live-cell microscopy-based assay that we previously described ^39^. Briefly, CL31 spheroids (from 100 cells each) were formed by overnight suspension culture, treated with either vehicle or recombinant human AREG (100 ng/ml) and clearance of a mesothelial monolayer was recorded over 24 hours. We found that AREG treatment significantly enhanced the mesothelial clearance ability of CL31 (Figure 7A-7B), suggesting that AREG may induce the phenotypic switch in CL31 by enhancing mesothelial clearance.

**Figure 7.**
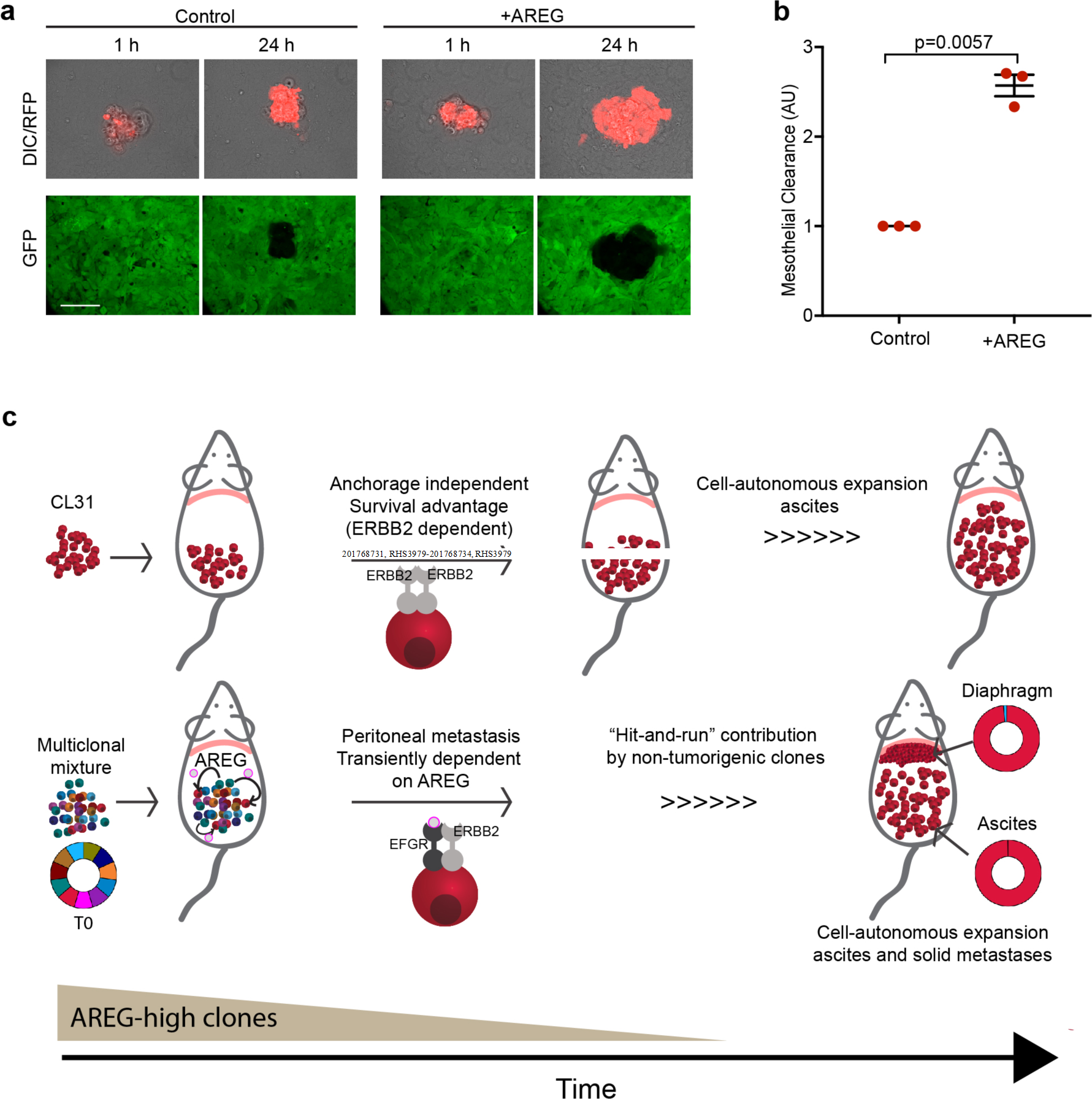
AREG enhances mesothelial clearance ability of CL31. (A) Representative differential interference contrast (DIC) and pseudocolored confocal fluorescence images of ability of vehicle-or AREG-treated CL31 cell clusters (red) to clear a mesothelial monolayer (green) at the indicated time points. Scale bar, 50μm. (B) Quantification of the mesothelial clearance area (black area within the green monolayer) cleared in 24 hours by CL31 spheroids treated with vehicle or AREG from three independent experiments. Mesothelial area cleared at the endpoint was normalized to the initial (1h) area of CL31 clusters (measured from the DIC images), as previously described ^39^. Relative clearance area of 20-30 clusters of CL31 of each condition per experiment were analyzed and averaged. Data shown as mean ± SEM of three independent experiments, shown in arbitrary units (AU), *P* value from Welch’s t test of the means. (C) Model illustrating the transient cooperative interactions involved in metastasis in this model system. CL31, which carries amplified ERBB2 and displays anchorage independent growth in vitro, forms only malignant ascites, but is unable to form solid peritoneal metastasis. Transient and non-tumorigenic AREG-high clones can act at an early temporal window, and are later dispensable, to induce CL31 solid peritoneal metastasis. AREG is required for metastasis of CL31, but not expansion in ascites or metastatic sites.

## Discussion

In this study, clonal populations of tumor cells were used as a model system to examine functional heterogeneity and spatio-temporal nature of interclonal interactions. Systematic comparison of the phenotypic properties and clonal growth dynamics of individual clonal populations as well as clonal mixtures revealed the importance of transient interclonal crosstalk between a tumor-initiating clonal population and ostensibly innocuous non-tumorigenic clones to strongly influence metastatic behavior. The mechanism involved in this interaction was elucidated by identification of unique molecular features of the clonal populations as well as genetic and non-genetic perturbations to define the critical regulators of this crosstalk. These studies highlight not only the importance of transient interactions of neoplastic cells in tumor progression, but also the value of such experimental platforms using clonal populations of tumor cells in order detect transient functional interactions inaccessible to analyses in bulk tumor cell populations.

Clonal populations with heterogeneous metastatic phenotypes have been reported in the triple negative breast cancer cell line MDA-MB-231 ^2, 3, 40, 41^. The findings from these studies show that each clonal population from this cell line expresses a set of intrinsic genes that determine its specific functional behavior, both with respect to metastatic ability, organ/tissue tropism, or responsiveness to the specific tissue environment ^42^. Findings from our model system provide evidence for an alternate, non-cell autonomous mechanism for acquisition of metastatic activity and support the previous proposed hypothesis that clonal subpopulations within a tumor cooperate with one another to promote metastasis ^11, 13, 43–46^. Importantly, our studies demonstrate that these cooperative interactions can be transient.

Ovarian tumor metastasis is believed to proceed through multiple steps, initially involving dissemination of tumor cells from the primary ovarian site into the peritoneum where there is selection for tumor cells that have acquired the ability to survive without cell-matrix interaction. Formation of solid peritoneal metastases requires additional steps, including intercalation into the mesothelial layer of peritoneal tissues and proliferation at those sites. The studies in this report are consistent with the model shown in Figure 7C. Based on both the low frequency of *ERBB2* amplification in the original tumor and the OCI-C5x parental cell line, we propose that acquisition of *ERBB2* amplification is a late event in the primary ovarian-localized tumor, generating a small subpopulation of cells with elevated ERBB2 expression which strongly enhances anchorage independence, thus promoting survival and proliferation of tumor cells shed into the peritoneum. Soluble factors, such as AREG, secreted by the non-tumorigenic and transient clonal populations that act in a “hit-and-run” fashion and are later dispensable, induce the ability of the *ERBB2*-amplified tumor cells to initiate solid peritoneal metastasis by promoting mesothelial clearance. AREG is not critical for growth or survival of the *ERBB2*-amplied tumors cells, but rather strongly promotes metastasis. It is possible that in absence of ligands, *ERBB2* amplification favors homodimerization of the ERBB2 receptors and thus maintains a ligand-independent and constitutively activated conformation that promotes anchorage independent proliferation and survival ^47^. When an EGFR associated ligand (e.g. AREG) is present, heterodimerization of EGFR-ERBB2 may promote prolonged and enhanced downstream signaling ^48, 49^ relative to that induced by ERBB2 homodimers. Studies have shown that AREG can promote invasion in multiple cancer models ^34, 50, 51^.

The interclonal interaction leading to acquisition of metastatic capability could take place either in the primary tumor site prior to dissemination, where the non-tumorigenic clones are able to survive because they are supported by the microenvironment and are not challenged under anchorage independent conditions in the ascites fluid, or within the peritoneum where possibly small clusters of heterogenous populations (ERBB2 amplified cells and AREG-high cells) are temporary coexisting and transiently support invasion of ERBB2-amplified cells and later die or are outcompeted. Our data demonstrate that AREG enhances mesothelial intercalation, providing one possible mechanism for the increased solid peritoneal metastases ^52^. This hypothesis would be consistent with the transient requirement for AREG and the absence of clonal populations other than CL31 in the solid peritoneal metastases. However, given that AREG treatment of CL31 was not sufficient to fully recapitulate the extent of solid peritoneal metastases of the parental or multiclonal mixtures and AREG blocking antibody did not completely eliminate the solid peritoneal metastases burden, it is likely that other secreted factors or additional receptor-ligand pairs are involved in the clonal cooperation. This is further supported by the finding that knockdown of ERBB2 protein in CL31 only partially impaired its anchorage independent growth and that only 18.9% on average of the cells comprising solid tumors generated by the parental line harbor *ERBB2* amplification, indicating that *ERBB2* amplification is not the sole mechanism of survival or metastasis in the parental line. It is likely that other clonal populations, which were not captured in our cloning process, are also able to metastasize.

There is evidence suggesting that co-existing heterogeneous tumor populations develop interdependencies since they support each other via symbiosis ^7–9^. Our findings suggest the existence of commensal relationships, in which one population (CL31) benefits from another without benefit to the latter ^43^. This implies that maintenance of tumor heterogeneity is not required for certain aspects of tumor progression, as temporal co-existence was sufficient to promote solid metastasis. Genetic analysis of the representation of clonal populations in primary versus secondary tumors has provided evidence that clonal populations are similar but not identical in primary and metastatic tumors ^53^ and conversely, that rare subclones in the primary tumor can give rise to metastases ^54, 55^. McPherson and colleagues ^56^ used mutational phylogeny analysis of autologous primary and distant peritoneal metastatic patient samples to show multiple modes of clonal spread in high grade serous ovarian carcinoma. While a few samples showed a high degree of polyclonal mixing and reseeding of multiple clones at distant sites, the majority of the clonal diversity emerged at the primary site followed by monoclonal seeding to distant intraperitoneal sites. The “hit-and-run” commensal model we propose can provide one explanation for how this pattern of metastasis can occur and could also help explain the dearth of metastatic drivers (or biomarkers) identified to date. These findings are of clinical relevance as the subpopulations providing these pro-metastatic factors would no longer be required or detectable at the distant metastatic sites. Further, clones with a low mutant allele frequency could be critical to drive or support metastasis. Identification of this type of commensal cooperative mechanism would not have been feasible with bulk analysis at static time points and at specific site. It is difficult to assess the generalizability of this mechanism of peritoneal metastasis in human cancer because static analyses of the clonal composition of matched primary and metastatic tumors fails to capture mechanisms involving unidirectional, transient interactions that drive metastasis but are dispensable for metastatic expansion.

Interestingly, our findings are consistent with models from evolutionary biology and cooperation theory models which show that cooperation in groups is often transitory and that cooperators can decrease in frequency within groups over time ^57, 58^. In the model presented here, clusters that include ‘cooperative’ AREG-producing cells (e.g. CL17) may allow more effectively clearance of the mesothelium in order to initiate metastasis. The evidence that these ‘cooperative’ cells could not be detected in metastases suggests that they are at a disadvantage and are evolutionarily outcompeted by CL31 cells within each cluster. Thus, Ovarian cancer metastasis offers an intriguing model for studying the evolutionary dynamics of cooperation among cancer cells.

The use of this model system revealed a critical role of transient and seemingly harmless clonal populations in tumor metastasis and uncovered an additional dimension of the complexity of metastasis-driving mechanisms. While an understanding of the impact of intratumoral heterogeneity on tumor behavior is emerging, our findings demonstrate that spatial and temporal aspects of clonal interactions must be taken into consideration.

## Materials and Methods

### Generation of clonal populations and virus production

OCI-C5x cell line was a kind gift from Dr. Tan Ince (University of Miami Miller School of Medicine, Ince et al., 2015). The cells were cultured and passaged in OCMI-L full medium (Live Tumor Culture Core, University of Miami) in a humidified incubator at 37°C with 5% CO2. OCI-C5x was engineered to co-express tdTomato and and *Gaussia* Luciferase via lentiviral transduction, as described above. After transduction, single cells positive for tdTomato were sorted under sterile conditions into 384-well plates containing OCI-C5x-conditioned media (50 µl/well) using fluorescence-assisted cell sorting (FACSAria cell sorter; BD Biosciences, Inc.). OCI-C5x conditioned media was prepared by seeding 7.5×10^5^ OCI-C5x cells in a 10cm plate with 10ml OCMI-L media and collected 48 hours after incubation. The media was spun down (3 minutes at 900 rpm) and the supernatant was stored at -20°C. Four hours after sorting, each well was visualized under a fluorescence microscope and only wells with one cell were included for our studies. Once cells reached confluence in the 384-well plate cells were trypsinized and replated into to larger wells with OCMI-L full media.

For barcode tagging of the clones, a lentivirus containing a unique barcode (Table S1) was introduced to a specific clone in an infection efficiency of 30%, and then selected for infected cells by 1μg puromycin for 4 days. Genomic DNA of each barcoded clonal population was isolated and equal amplification rate among all population was confirmed.

### Plasmids, shRNAs, and Virus Production

CSCW-GLuc-IRES-CFP was a generous gift from Dr. Bakhos Tannous lab ^23^. To replace the CFP with tdTomato, we transferred the Kpn1-digested fragment from pBS-IRES-tdTomato-WRE plasmid into the Kpn1-digested CSCW-GLuc-IRES-CFP plasmid followed by standard quick ligation protocol. Overexpression and knockdown of specific target genes in these cells was performed via lentiviral transduction using standard protocols, following selection with 1μg/ml Puromycin for 4 days. Flag-HA-AREG and Flag-HA-GFP were a generous gift of the Harper W. lab (Harvard medical school, Boston, MA). shRNA vectors for ERBB2 were purchased from Dharmacon (RHS3979-201785743, RHS3979-201768731, RHS3979-201768734, RHS3979-201768735). shRNA-GFP control (Addgene, #30323).

### Analysis of barcode distributions

The *in vitro* multiclonal mixture cultures were sampled each time they were passaged, i.e. approximately 2 x 10^6^ cells were collected following cell counting by centrifugation, and stored at -80°C. Tumor tissue samples were also stored at -80°C prior to genomic DNA isolation. Tumors were mechanically homogenized on ice and entire tumor tissue was transferred to an Eppendorf tube for genomic DNA isolation. The genomic DNA was isolated using the QIAamp DNA Mini Kit (Qiagen, Cat. No. 51306) and DNA concentrations were measured on the Epoch microplate spectrophotometer (BioTek). The barcodes were subsequently isolated by PCR on 3 µg genomic DNA per sample (max. 1 µg per PCR) using a common forward primer and set of reverse primers with unique index sequences that allow for multiplexing of samples. Per sample, PCR products were combined and isolated using the QiaQuick PCR purification kit (Qiagen, Cat. No. 28106). The quality and concentration of the PCR products was determined using a 2200 TapeStation and D1000 screen tapes (Agilent Technologies). To generate the NGS libraries, samples were mixed in equal proportions. The libraries were isolated from a 2% agarose gel using the QiaQuick gel extraction kit (Qiagen, Cat. No. 28706) and their quality and concentration were determined using a 2200 TapeStation and D1000 screen tapes (Agilent Technologies), and q-PCR. Each library was sequenced on a MiSeq (Illumina) by the Biopolymers Facility at Harvard Medical School using the following primers:

WSL_NGS_Barcode_Seq: GCGACCACCGAGATCTACACACTGACTGCAGTCTGAGTCTGACAG

WSL_NGS_Index_Seq: GATCGGAAGAGCACACGTCTGAACTCCAGTCAC

Index sequences were used to demultiplex the samples. Barcode sequences that matched the 15xWS design, a Phred quality score of 10 or greater for each position and an average Phred quality score greater than 30, were selected for further analysis. First, we selected the barcode sequences that were used in the experiment and then we normalized the barcode counts to their mean fractions in the T=0 reference samples.

For each sample, the barcode fractions were computed by dividing the number of reads of a given barcode by the total number of reads for all barcodes in that sample (Supplementary spreadsheet 1, Raw barcode counts).

### Doubling time

Each population was plated into a 6-well multiwell plate (Corning) at a density of 7×10^3^ cells. Cells were harvested by trypsinization on days 1, 3 and 5 and counted using a particle counter (Z1; Beckman Coulter, Inc.). Experiments were carried out in triplicate. Doubling time was calculated with the assumption of exponential growth. The number of generations was calculated using the following formula: [(*log*(*Day* 1) − *log*(*Day* 5))/ *log* (2)]. Doubling time was derived by dividing the duration of the experiment (96 hours) by the number of generations.

### Soft agar assay

The soft agar assay was performed in 6-well plates, the assay was carried out in duplicates for each tested group. For each tested group 2×10^4^ cells were added to 1 ml of 0.4% low-melt agarose solution (Sigma) in OCMI-L media and transferred to a well with a bed of growth media with 0.5% low-melt agarose. At day 21, viable colonies were stained with iodonitrotrazolium chloride (Sigma) (25 mg/ml) (Sigma-Aldrich).

Subsequently colonies were imaged with a dissecting microscope (2.5x). ImageJ was utilized to count & analyze particles. Data was analyzed GraphPad Prism (GraphPad, Inc.). Images shown are representative of at least three independent experiments.

### Copy Number Analysis

Genomic DNA was extracted from each sample using the DNeasy Blood & Tissue Kit (Qiagen, 69504), followed by acoustic sonication using Covaris S220 sonicator. DNA concentration was quantified using the BioTek microplate reader (BioTek Instruments, Inc.). Next, the DNA was sparse-sequenced to infer the copy number profiles. The sequencing libraries were constructed by adding 3’ adenylation and ligating barcoded sequencing adaptors to the fragmented DNA using NEBNext DNA Library Prep Master Mix set for Illumina (NEB, E5040L). The generation of barcoded sequencing adapters can be found in ^59^. The barcoded libraries were pooled and sequenced on a sequencing lane of Illumina HiSeq 2500.

The sequencing data was analyzed following the informatics procedure in Baslan *et al* ^60^. Briefly, the sequencing reads were mapped to the human assembly hg19 using bowtie 2 and the Genome Analysis Toolkit (GATK) was used to locally realign the BAM files at interval that had INDEL mismatches before PCR duplicate marking with Picard. Copy number was calculated from read density by dividing the genome into variable bins and counting the number of unique reads in each 200Kb interval. Bins at the centromeric and telomeric regions were filtered to remove false-positive errors. We then applied Loess normalization to correct for GC bias ^60^. The copy number profiles were segmented using circular binary segmentation as in Olshen *et al*. 2004 ^61^.

### Whole-Exome Sequencing and Mutation Calling

Whole-exome libraries were prepared from DNA extracted from the individual clones, parental line, and the patient’s blood. DNA was sonicated to 150bp using the Covaris e220. Libraries were created using either the Agilent SureSelect XT protocol, or the Kapa HyperPrep protocol (Kapa Biosystems). Hybrid capture was conducted for 24hrs using the Agilent human all exon v4 + UTR capture baits. For two samples (Patient blood and parental line) an additional library was created using the Kapa HyperPrep protocol and captured using the Agilent human all exon v5 baits. Libraries were 100 bp paired-end sequenced on a Illumina HiSeq 2000.

Reads were aligned to hg19 using BWA mem, local realignment and quality score recalibration were conducted using GATK, PCR duplicates were marked using Picard Tools. Somatic mutation calls were made using Mutect (v 1.1.7), patient blood was used as the match normal, calls were filtered against the dbSNP database (v 137) and rescued if they were present in the COSMIC (v 54) databases. As a final filter, mutation calls were required to have 30X depth in the tumor, 10X in the normal, at least a 10% mutant allele frequency in the tumor and less than 2% in the normal. After the full list of filter-passing mutations was generated, samtools mpileup was generated for each mutation site in each of the clones to look for the presence of the mutant reads below our initial calling threshold. Functional annotation of mutation calls were conducted using Oncotator (http://portals.broadinstitute.org/oncotator/).

Mutational signature analysis was conducted using a list of all filter-passing mutations for each sample (Supplemental spreadsheet 2, all mutants tab). The PCAWG trinucleotide signatures were used for cosine similarity signature assignment using the mutational patterns R package. Signatures were filtered to only include those contributing at least 200 total mutations across all samples (https://github.com/UMCUGenetics/MutationalPatterns).

### Tree Building

Whole-exome mutations were filtered to include only those present at 30X depth across all samples, and present at a VAF >=0.25 in at least one sample. The 7,402 resulting mutations were the classified as present in a sample (VAF >= 0.25), absent (VAF < 0.10), or ambiguous (0.10 <= VAF < 0.25). The resulting matrix was then input into mpboot to generate a maximum parsimony tree using 1,000 bootstrap iterations ^62^. All nodes had bootstrap support of 1.0. For the mutations composing each branch, the mean VAF of those mutations in the parental clone was determined, and the branch colored accordingly.

### RNA Sequencing

Total RNA was extracted from each sample using the RNeasy Mini Kit according to manufacturer’s protocol (Qiagen, Inc.). The RNA concentration was measured using the Epoch microplate spectrophotometer (BioTek). Prior to library generation for RNA-sequencing, the quality of the RNA was determined on a 2200 TapeStation analyzer using RNA screen tapes (Agilent Technologies). The mRNA libraries were generated by the Biopolymers Facility at Harvard Medical School (http://genome.med.harvard.edu, Boston, MA) and included poly-A enrichment and the Directional RNASeq Wafergen services. The quality of these libraries was assessed on a 2200 TapeStation analyzer using D1000 screen tapes (Agilent Technologies), and by qPCR. In total, 12 mRNA libraries (one sample for the OCI-C5x parental cell line, and one sample of each clonal population) were multiplexed into one pool and sequenced on Illumina HiSeq2000 instrument (Illumina Inc., San Diego, CA). Adapter sequence was trimmed using Trimmomatic-0.30. Trimmed reads were mapped with bowtie2-2.1.0 to the hg19 version of the human genome. The HTSeq-0.5.4 and EdgeR packages were used to quantify counts per gene and perform differential expression analysis, respectively.

Growth factors/cytokines and associated receptors were identified from AmiGO 2 (http://amigo.geneontology.org/amigo) and the Database of Interacting Proteins (https://dip.doe-mbi.ucla.edu/dip/DLRP.cgi). Hierarchical clustering was performed in R 3.5.1 using Spearman correlation and average linkage. Heatmaps were generated with the heatmap.2 package. RNA-seq data has been submitted to the Gene Expression Omnibus (GEO), GSE123426.

### Animals and *In Vivo* Procedures

#### Xenograft studies

All animal studies were performed according to protocols approved by the Harvard University’s Institutional Animal Care and Use Committee (IACUC). Eight-week old female NOD-*scid* IL2Rgamma^null^ (NSG) mice were purchased from the Jackson Laboratory (Bar Harbor, Maine). For each experiment, Gluc-tdTomato expressing cells were injected intraperitoneally to 8-10 weeks old female NSG mice.

The mice were euthanized 10 weeks after tumor cell injection. Each mouse was dissected and visually inspected for solid tumor metastasis under a fluorescence dissecting microscope for tdTomato signal, organs with tumors were collected, fixed with 10% formalin (Westnet-Simport), paraffin embedded and sectioned for H&E staining. Peritoneal cavity was washed with 1x PBS to collect malignant ascites into graduated 15 ml conical tubes, then the cells were spun down and the volume of cellular pellet was assessed by the volume marks on the tube since precise measurement of the pellet volume is difficult due to its viscosity. An aliquot of the malignant ascites for each sample was mixed with equal volume of histogel (VWR), incubated for 20 minutes in room temperature to solidify, fixed in 10% formalin followed by paraffin embedding, sectioning and H&E staining. Three to five mice per group were used for each experiment and at least two independent experiments were performed.

#### Solid metastases tumor burden scoring

On completion of immunohistochemical staining of the tissue samples, a pathologist examined the tissue slides in a blinded manner and documented the tumor burden. The classifications of tumor burden were based on a five-point scale: 0, no tumor cells; 1, few tumor cells; 2, few and small clusters of tumor cells; 3, large clusters of tumor cells, deposited along large area of the diaphragm; 4 and 5, bulky tumor deposits on part or the entire diaphragm, categorized as either 4 or 5 depending on the size of the deposits within this group. Two slides were scored for each sample and the average of the score was calculated.

#### Solid metastases tumor burden analysis by weight

Tumor masses identified on the diaphragm were cut out from the diaphragm and weighed. Tumor weight, in addition to the tumor burden scoring method, was used to monitor effect of amphiregulin blocking antibody on tumor burden since in this specific experimental design both the control and the tested groups had weighable tumors for comparison.

#### *In vitro* Gaussia luciferase (G-Luc) activity

Blood from the submandibular vein of each mouse was collected into tubes with Heparin (Henry Schein animal health supply, Ohio) at 24 hours after tumor cell injection and every week thereafter for up to 10 weeks. Five microliters of blood from untreated mice (negative control) and of treated mice were transferred into each well of white 96-well plate in duplicate. To test the G-Luc activity in blood, coelentrazine solution (Prolume LTD, AZ) at a final concentration of 20μM was prepared with DMEM media (Life technologies) and incubated for 30 minutes at room temperature, then 50μl of the coelentrazine was added using multichannel pipettor to each well and bioluminescence signal was measured immediately using a plate luminometer (MLX luminometer, Dynex Technologies, Chantilly, VA). Since G-Luc is not equally expressed across all clonal populations and G-Luc values can vary between experiments, fold-change in G-Luc activity was calculated by normalizing to Day 1 G-Luc levels (24 hours after injection) to allow better comparison between clones and across multiple experiments. G-Luc fold-change for each mouse was averaged for each time point and graphed on a log scale using GraphPad Prism.

#### Amphiregulin blocking antibody *in vivo*

A total of 3×10^6^ cells of an equal mixture of CL31 and CL17 were injected I.P. to 8-10 week old female NSG mice. Immediately after cell injection, mice were treated with 200μg monoclonal AREG blocking antibody (AREG 37.4 Sigma-Aldrich, Israel) in a 200μl PBS (I.P. injection), twice a week for 10 weeks. Control group was treated with PBS vehicle control. Tumor growth dynamics was monitored by measuring G-luc activity from blood and tissues were collected as mentioned above. Solid masses on the diaphragm were collected and weighed.

#### Human recombinant AREG peptide *in vivo*

A total of 3×10^6^ cells were injected I.P. into 8-10 week old female NSG mice. Immediately after cell injection, mice were treated with human recombinant Amphiregulin (Cat# 100-55B, Peprotech) dissolved in 0.1% BSA (Sigma) to a final concentration of 1μg/μl. Mice were treated by I.P. injection with either 5ug/mouse AREG in 200μl sterile water (Gibco) or BSA vehicle control (0.0025% BSA final concentration) as follows: starting on the day of cell inoculation and every day for a week, then every other day on the second week and twice a week on the third week. No AREG supplementation from week 4 through week 10.

### FISH analysis

Tumor blocks of FFPE were sectioned (5 µm in thickness). For each specimen, one sectioned slide was stained with H&E and used by rodent pathologists to mark tumor areas. For the original patient tumor sample, the tumor area was identified by pathologist and by S100A1 staining that differentiates carcinoma cells from normal epithelial cells in the ovary. The sections were delivered to the Cytogenomics Core laboratory (cytogenomics.bwh.harvard.edu) at the Brigham and Women’s Hospital (Boston, MA) for HER2 following manufacturer protocol of a commercial two-color FISH probe set (PathVysion, Abbott Molecular). For each sample 60-100 cells were scored for ERBB2:CEP17 ratio. Each sample was scored independently by two technicians.

### Immunohistochemistry Staining

Formalin-fixed tumor samples derived from mouse xenografts were processed and embedded in paraffin. Tissue sections were deparaffinized and antigen retrieval was achieved by use of heat-induced epitope retrieval with pH 6.0 citrate buffer (Sigma). Tissue sections were stained with H&E, anti-ERBB2 antibody (Dako, A0485), or anti-S100A1antibody (Dako, Z031129-2). Immunostained slides were counterstained with hematoxylin (Sigma). Images of tissue sections were captured at 10x magnification to create a merged image of the entire section using Olympus VS120 Slide Scanner.

### Secreted Amphiregulin Measurement

OCI-C5x parental cells and each individual clone were seeded at the same density (7.5·10^5^ cells in a 10 cm plate) with 5 ml of 199:MCDB105 media with 2% heat inactivated fetal bovine serum and incubated for 48 hours. The media was collected and spun down (3 minutes at 900 rpm) and the supernatant was used or alternatively, kept at -20°C. ELISA test for AREG levels was performed using 100ul of each sample in duplicates and following the manufacturer protocol (Abcam, Ab99975).

### Mesothelial clearance assay

Comparable size multicellular spheroids of CL31 were prepared by culturing 100 cells of CL31 per well on low attachment 96-well round bottom plates (Westnet, 7007), cells were cultured in a media of equal mix of 199 media (Gibco) and MCDB105 media (Sigma) (199:MCDB105 media) with 2%FBS and incubated overnight at 37°C to promote spheroid formation. Concurrently, 250,000 ZT mesothelial cells per well were plated on fibronectin-coated (5 μg/ml, Sigma-Aldrich) 24-well glass-bottom culture dishes (MatTek corporation, P24G-1.5-13-F). The mesothelial cells were incubated overnight at 37°C to form confluent monolayers. After the 24-hour incubations, 10-15 multicellular spheroids of each condition were transferred to the wells containing the mesothelial monolayers in duplicates wells. The spheres were allowed to settle for two hours prior to starting live imaging. Imaging was performed using a Nikon Spinning disck confocal microscope with the integrated Perfect Focus System and low (×20, 0.75 NA) magnification/NA DIC optics, Nikon fast (<100-ms switching time) excitation and emission filter wheels, Nikon linear-encoded motorized stage, Hamamatsu ORCA-AG cooled CCD camera, custom-built microscope incubation chamber with temperature and CO2 control, Nikon NIS Elements AR software v3, and TMC vibration-isolation table. Over 20 spheroids were imaged per condition. Phase-contrast, GFP and RFP images were captured every 20 minutes for 22 hours. To quantify the mesothelial clearance area, the non-fluorescent area, created by the invading spheroid, in the GFP mesothelial monolayer images was measured at 22 hours and divided by the initial area of the cancer spheroid at time 0. All measurements were taken using ImageJ software.

### Quantitative PCR

mRNA prepared from cell extracts using the RNeasy Mini Kit (QIAGEN) was reverse-transcribed into cDNA using the qScript cDNA synthesis kit (Quanta Biosciences). Real-time PCR was performed on an ABI PRISM 7900HT or QuantStudio 7 Flex Real-Time PCR System (Life Technologies) with Power SYBR Green PCR Mix (Life Technologies). Amphiregulin forward primer: TGATCCTCACAGCTGTTGCT; Reverse primer: TCCATTCTCTTGTCGAAGTTTCT; RPLPO forward primer: ACGGGTACAAACGAGTCCTG; Reverse primer: CGACTCTTCCTTGGCTTCAA

### Immunoblot Assay

Samples were lysed in cell RIPA lysis buffer (Boston Bioproducts) supplemented with protease and phosphatase inhibitor cocktails (Roche life science). Protein concentrations were determined by BCA protein assay (Life Technologies). Equal protein amounts were denatured in SDS buffer (1% SDS, 8% Glycerol,62.5mM Tris-HCL, pH 6.8) with 1.5% β-mercaptoethanol for 5 minutes. Protein were analyzed by SDS-PAGE and immunoblot using indicated antibody. Blots were imaged with Odyssey CLx infrared imaging system (LI-COR) and are representative of at least two independent experiments.

### Statistics

The specific statistical tests used and resulting p values are indicated in the figure legends. Unless indicated otherwise, all statistical tests were carried out using Graphpad Prism version 7.0. Statistical analysis for the tumor burden scoring was performed using R software.

## Acknowledgments

We thank the Brugge lab members for helpful discussions, Dr. Angie Martinez Gakidis for scientific editing and critical review of the manuscript, Dr. Bakhos A. Tannous for the kind gift of the secreted Gaussia Luciferase construct, Dr. Laura Pontano Vaites from the Dr. Wade J. Harper lab for providing the amphiregulin lentiviral vector, Dr. Hyo-eun C. Bhang and Dr. Frank Stegmeier for providing the barcode constructs, Min Hu from Dr. Nicholas Navin’s lab for assisting with the CNA analysis, Anita L. Hawkins and Shumei Wang for their help with the ERBB2 FISH analysis. The time lapse movies were acquired with the support of The Nikon Imaging Center at Harvard Medical School. Whole-slide scans of tissue were acquired at the Neurobiology Imaging Facility at Harvard Medical School. The assay for Luciferase activity was performed at ICCB-Longwood Screening Facility at Harvard Medical School.

## Author Contributions

S.N.A. and J.S.B. initiated the study and conceived the project, designed experiments, interpreted results, and wrote the manuscript. S.N.A. performed most of the experiments. H.J.K. optimized and performed the barcode representation analysis. T.B. and P.S. carried out the whole exsome sequencing and analysis. M.L and N.N carried out the CNA analysis. C.T.L. generated the tdTomato-Gluc vector. L.L. assisted with the xenograft model. T.K, C.H. and K.S.G. assisted with the soft agar colony formation and mesothelial clearance assays. R.F. and B.R.R generated the original PDX model. T.I generated the OCI-C5x cell line. L.M.S performed bioinformatics analysis. R.T.B. analyzed the histological sections and blindly scored tumor burden. G.M. participated in experimental design and interpretation. A.A. contributed a section to the discussion about evolutionary biology and cooperation theory.

All authors participated in manuscript revisions.

## Financial support

This research was supported by NCI grant CA181543 (Joan S. Brugge), R33CA214310 and OC130649 (Tan A. Ince), The Knight Cancer Institute (Paul Spellman), 5T32GM07133 (Timothy Butler) and 129098-RSG-16-092-01-TBG (Nicholas Navin).

## Disclosure of potential conflicts of interest

The authors declare no potential conflicts of interest.

