## Supplementary Figures for "Transient Commensal Clonal Interactions Can Drive Tumor Metastasis"

**Supplementary Data:**

**Figure S1.**

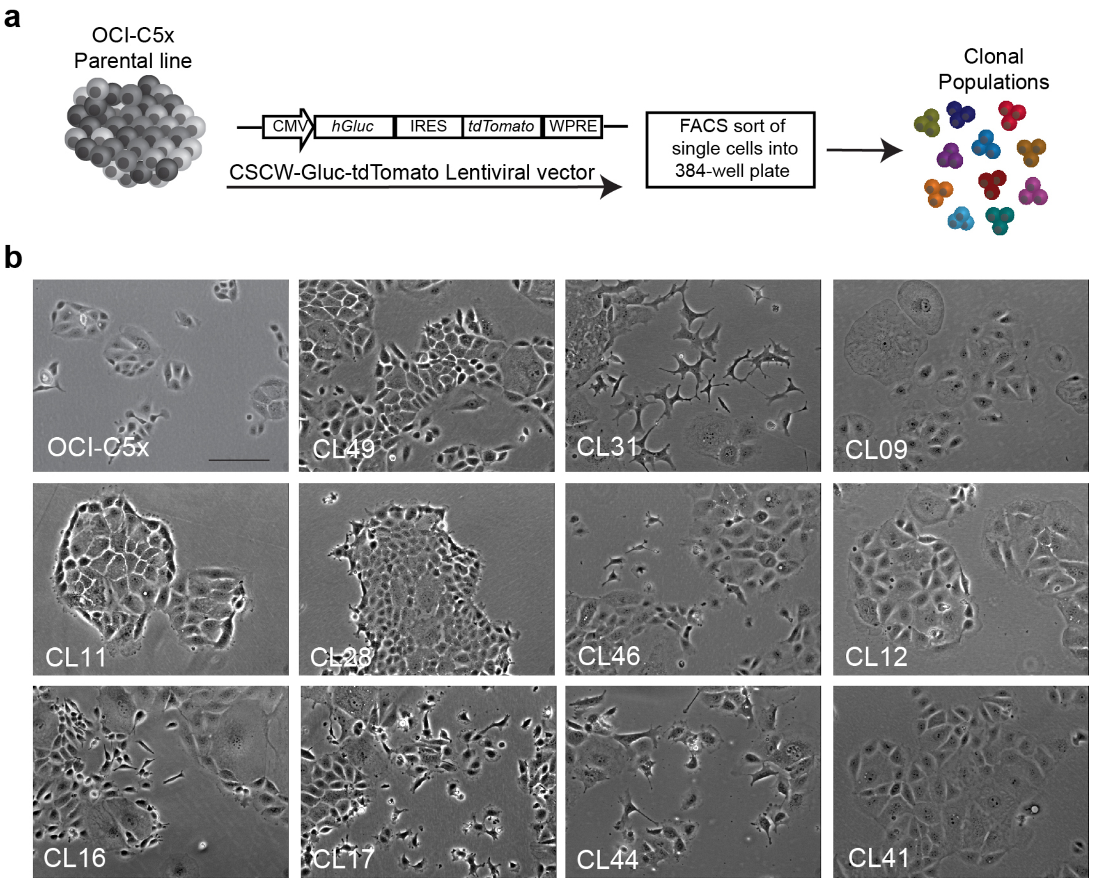


**Figure S1. Clonal populations derived from OCI-C5x cell line vary in morphology.**

(A) Schematic description of generation of the clonal populations from OCI-C5x. (B) Representative phase contrast images of parental OCI-C5x line and clonal populations. Scale bar, 100μm.

**Figure S2.**

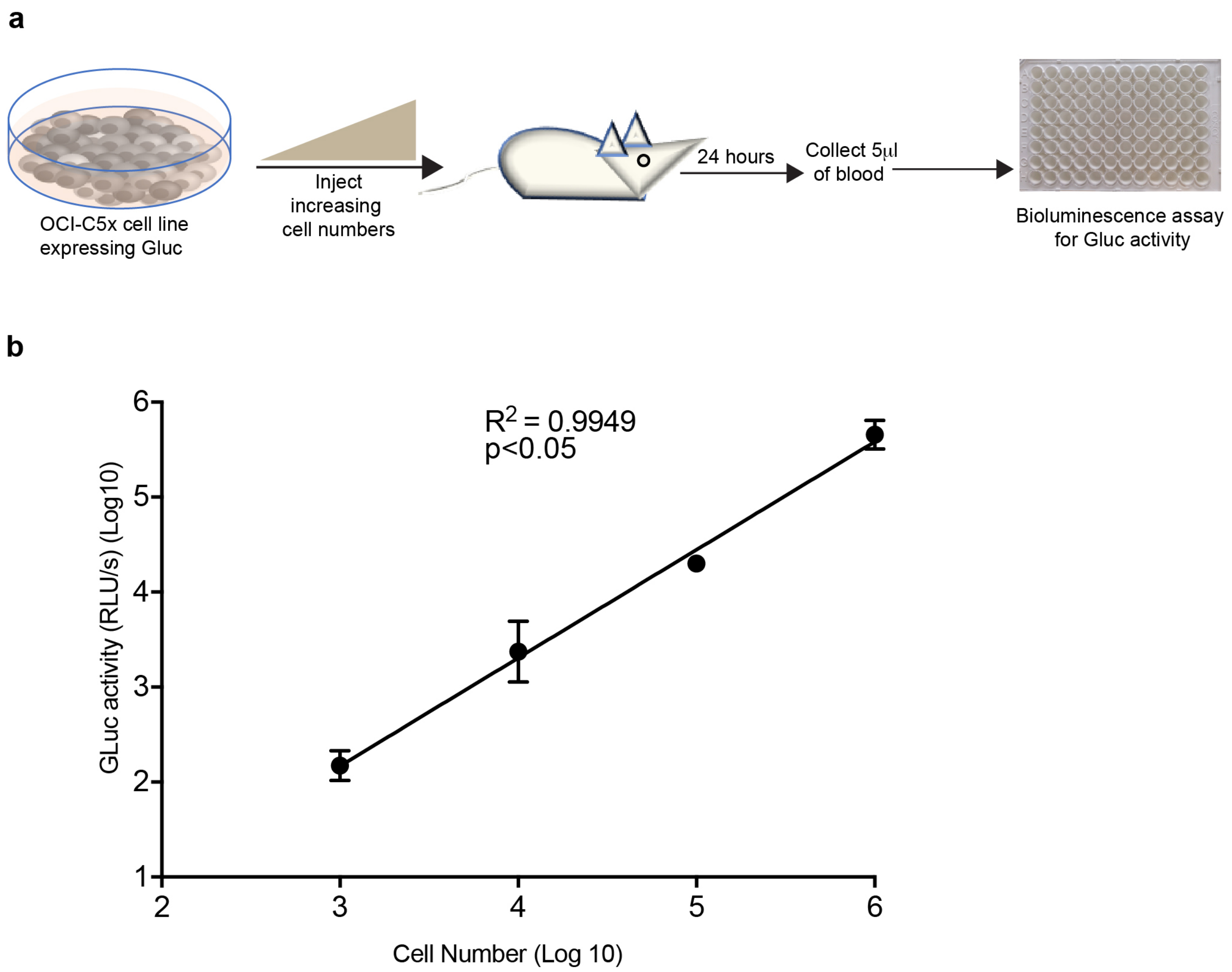


**Figure S2. Measurement of tumor growth dynamics by luciferase assay.**

(A) Increasing number of OCI-C5x cells expressing Gaussia luciferase (Gluc) were injected intraperitoneally into NSG mice. Blood samples were collected 24 hours post injection and Gluc activity was measured on 5μl of blood in duplicates. Values are the average of the duplicates. (B) Linear regression of number of injected cells with *ex-vivo* luciferase activity in blood detected at 24 hours post injection. $p$ <0.05, two-tailed t test.

**Figure S3.**

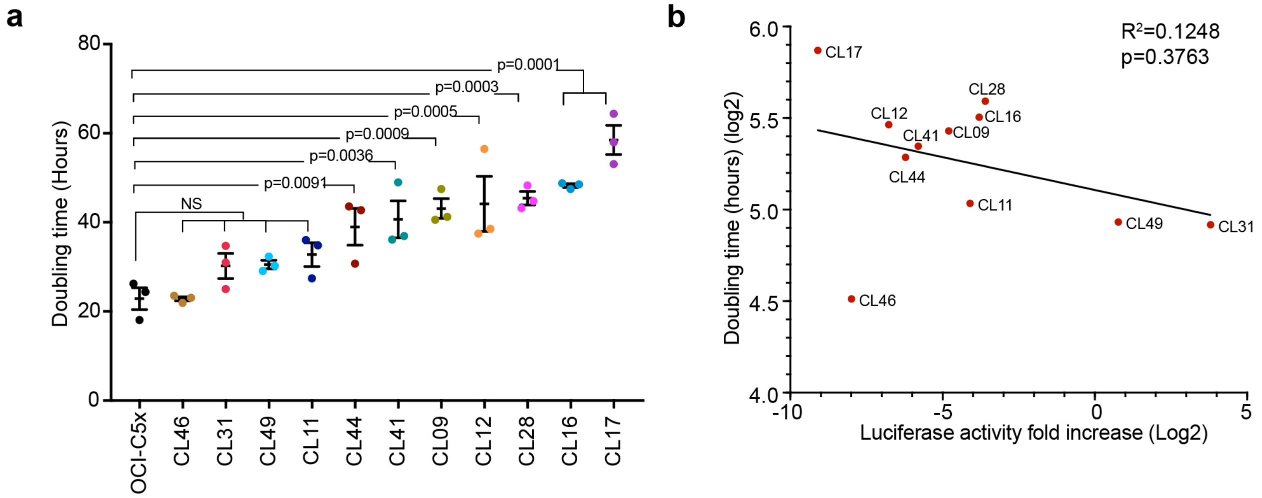


**Figure S3. Clonal populations derived from OCI-C5x cell line vary in population doubling time.**

(A) Doubling time of OCI-C5x and clonal lines grown in triplicate and for over 5 days. Doubling time was computed as follows: 24*[(LOG (cell number day 1)-LOG (cell number day 5))/LOG (2)]. Data shown as mean± SEM from three independent experiments. *p* values from one-way ANOVA, corrected for multiple comparisons using Dunnett's method. (B) Linear regression analysis of doubling time and tumor burden (measured by luciferase activity in blood samples collected at 10 week end point) for all 11 clonal populations.

**Figure S4.**

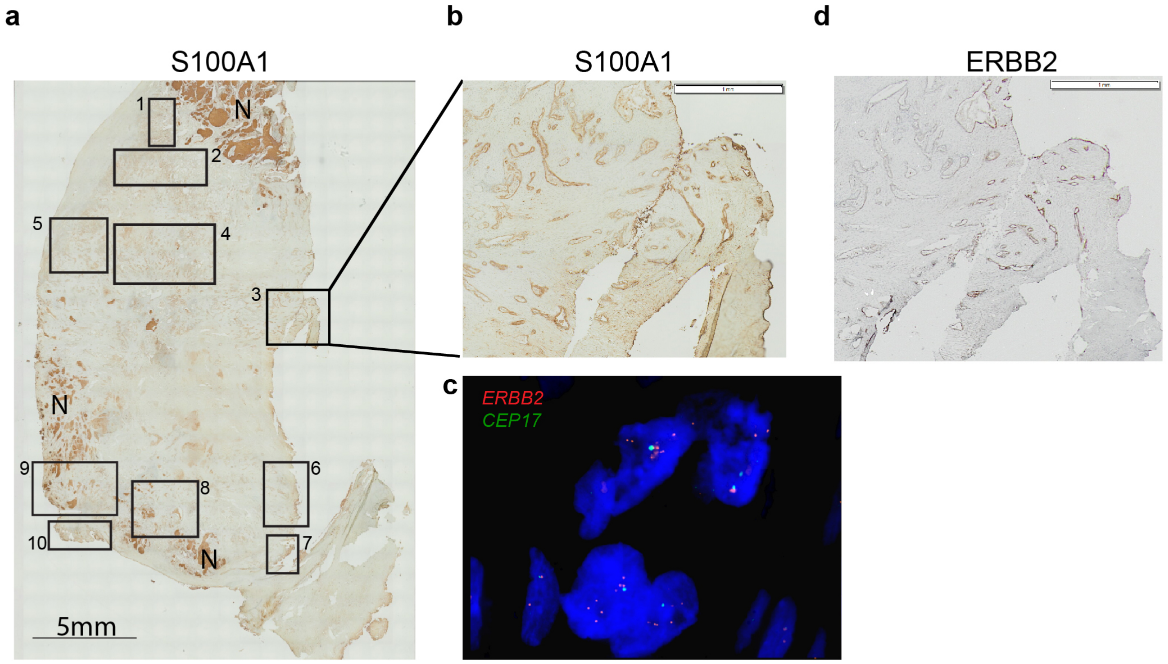


e

| Region # / *ERBB2*:CEP17 score | 1 | 2 | 3 | 4 | 5 | 6 | 7 | 8 | 9 | 10 |
|---|---|---|---|---|---|---|---|---|---|---|
| % of scored cells with *ERBB2*:CEP17≥2 | 0 | 1.66 | 26 | 23.3 | 13.3 | 25 | 16 | 10 | 20 | 0 |
| Average of *ERBB2*:CEP17 ratio from cells above | NA | 2 | 2.96 | 2.42 | 2.6 | 2.7 | 2.46 | 2 | 2.5 | NA |

**Figure S4. Identification of *ERBB2*-amplified cells in the original patient sample from which OCI-C5x was derived.**

(A) Histology section of the patient ovary (primary site) from which OCI-C5x was derived, stained for S100A1 to identify ovarian tumor cells from normal ovarian epithelial cells. Regions marked with "N" indicate necrotic areas that are false positive for S100A1 staining. The boxes indicate the ten regions chosen for *ERBB2*:CEP17 scoring by FISH. Scale bar 5mm. (B) Higher magnification of Region #3 (S100A1). Scale bar 1mm

(C) Representative image of FISH staining for *ERBB2* (red) and CEP17 (green). (D) IHC of ERBB2 staining of Region #3. Scale bar 1mm. (E) Summary table of the *ERBB2*:CEP17 patterns in all the tested regions.  Percentage of scored cells with *ERBB2*:CEP17 equal or greater than two are shown and *ERBB2*:CEP17 signal ratio was calculated for this group of cells. Cells with only one CEP17 signal and two ERBB2 signals were excluded from this analysis to reduce false positives.

**Figure S5.**

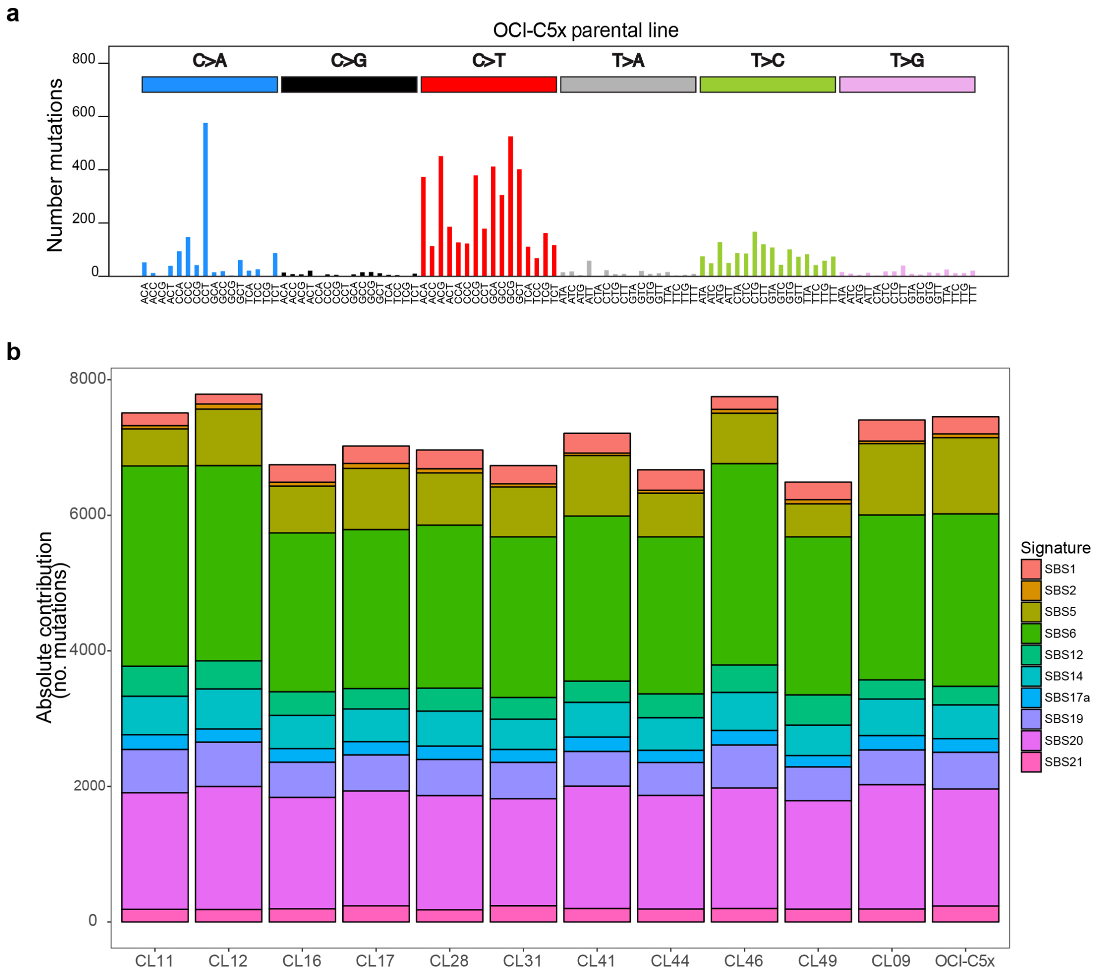


**Figure S5. Mutational signatures and allele frequency analyses.**

(A) Spectrum of 96 trinucleotide mutational contexts for 7,484 filter-passing mutations in the parental line. The six substitution types are shown above, and the bases immediately 5' and 3' to the mutated base are shown below. (B) Comparison of Single Base Substitution (SBS) mutational signatures present in each clone. No significant differences between the samples.

**Figure S6.**

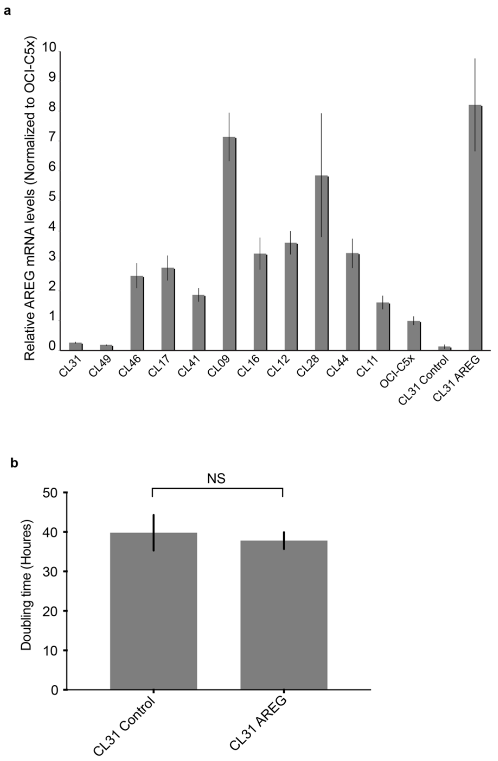


**Figure S6. AREG overexpression in CL31.**

(A) qRT-PCR measurements of mRNA levels of AREG in the OCI-C5x parental line, the clonal populations and CL31 transduced with empty vector (Control) or AREG. Measurements from three technical replicates were normalized to *RPLPO* mRNA levels and expressed as fold change compared to the OCI-C5x parental line. (B) Doubling time of CL31-Control and CL31-AREG lines grown in triplicate and for over 5 days. Doubling time was computed as follows: 24*[(LOG (cell number day 1)-LOG (cell number day 5))/LOG(2)]. Data shown as mean ± SEM of three independent experiments, *P* value from Welch's t test of the means. NS: not significant.

**Figure S7.**

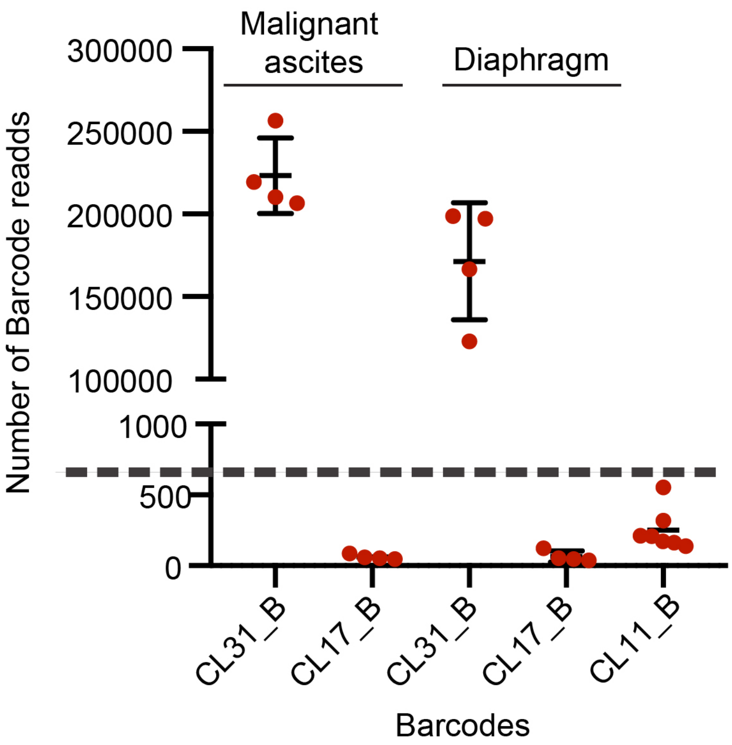


**Figure S7. Malignant ascites and solid peritoneal metastases generated by the CL31:CL17 mixture consist exclusively of CL31.**

(A) CL31 and CL17 barcodes counts (CL31_B, CL17_B, respectively) of indicated samples collected from mice at 10 weeks following IP injection of CL31:CL17 mixture. The barcode count for CL11, which was not present in the mixture, was used as a negative control to set the noise threshold of the barcode counts (dashed line).

**Table S1: Barcodes ID and their target clones**

| Clone_ID | Barcode_ID | Barcode Sequence |
|---|---|---|
| CL09 | WS-01-SingleWS | TGACTGTGAGTGTCTGTCACAGTGTGTGAG |
| CL44 | WS-08-SingleWS | TGTGTCTGACTGACTGACAGTGACACACTG |
| CL49 | WS-21-SingleWS | TCTGTGTCACACACTCTGTCTGAGAGTGTC |
| CL17 | WS-17-SingleWS | TGACTCAGAGAGTCTCTGTGTCTGTCAGAC |
| CL41 | WS-23-SingleWS | ACTGTCTGAGACAGAGAGTGTGACAGTCAG |
| CL16 | WS-16-SingleWS | TCTCTGAGACACAGTCAGAGTCACAGTGTG |
| CL46 | WS-19-SingleWS | AGACAGACTCTCAGTCTGTCAGACAGTGAG |
| CL31 | WS-14-SingleWS | AGTGTCACTGTGTGACTGAGAGTCTGACAG |
| CL11 | WS-03-SingleWS | TCTGACACTCAGACTCAGTGACTGTGACTG |
| CL28 | WS-13-SingleWS | TCAGTCTCAGTCTCACTGTGTGTCACTCTC |
| CL12 | WS-04-SingleWS | TGACAGAGTGTGTCAGAGTGTGAGTGAGTG |
